# Single-Cell Resolution of Lineage Trajectories in the Arabidopsis Stomatal Lineage and Developing Leaf

**DOI:** 10.1101/2020.09.08.288498

**Authors:** Camila B. Lopez-Anido, Anne Vatén, Nicole K. Smoot, Nidhi Sharma, Victoria Guo, Yan Gong, M. Ximena Anleu Gil, Annika K. Weimer, Dominique C. Bergmann

**Affiliations:** Department of Biology, Stanford University, Stanford, CA 94305-5020, USA; Howard Hughes Medical Institute, Stanford University, Stanford, CA 94305-5020, USA; Organismal and Evolutionary Biology Research Programme, Biological and Environmental Sciences, University of Helsinki, Helsinki 00014, Finland; Genentech Inc, 1 DNA Way, South San Francisco, CA 94080; Department of Plant Biology, 2203 Life Sciences Building, University of California, Davis, CA 95616, USA; Department of Genetics, Stanford University School of Medicine, Stanford, California 94305, USA

## Abstract

Dynamic cell states underlie flexible developmental programs, such as with the stomatal lineage of the Arabidopsis epidermis. Initial stages of the lineage feature asynchronous and indeterminate divisions modulated by environmental cues, enabling cell fate flexibility to generate the requisite density and pattern of stomata for a given environment. It remains unclear, however, how flexibility of cell fates is controlled. Here, we uncovered distinct models of cell state differentiation within Arabidopsis leaf tissue by leveraging single-cell transcriptomics and molecular genetics. Our findings resolved underlying heterogeneity within cell states of the flexible epidermal stomatal lineage, which appear to exist along a continuum, with progressive cell specification. Beyond the early stages of the lineage, we discovered that the core transcriptional regulator SPEECHLESS is required for cell fate commitment to yield stomatal guard cells. Overall, our work has refined the stomatal lineage paradigm and uncovered progressive cell state decisions along lineage trajectories in developing leaves.

## INTRODUCTION

Cellular fate specification and differentiation are core features of developmental programs in multicellular organisms. Molecular genetics established the classical view of cell fate commitments as discrete and sequential stages, but recent technological advances with single-cell RNA sequencing (scRNA-seq) have revealed ways in which cell state decisions are continuous and heterogeneous. Cell states may indeed be now defined by gene expression profiles that mark states as putatively dynamic instances of a given cell transition or identity, corresponding to cell function and lineage relationships (Morris, 2019; Sagar and Grün, 2020; Wagner et al., 2016). While remarkable progress has been made to elucidate cell states within a range of tissue and animal systems (Karaiskos et al., 2017; Schaum et al., 2018; Sebé-Pedrós et al., 2018; Siebert et al., 2019; Tintori et al., 2016; Wagner et al., 2018), much is to be learned from state dynamics in plants. In addition to exhibiting many differentiated cell types with no counterparts in animals, plants display extreme biological flexibility, longevity, and regenerative capacity. Multicellularity, and therefore developmental processes, arose independently in plants and animals, though there are many cases of convergence in the deployment of specific molecular regulators and developmental strategies.

Recent applications of scRNA-seq to plants include detailed analyses of regeneration and meiosis utilizing hundreds of highly selected cells (Efroni et al., 2016; Nelms and Walbot, 2019). Conversely, atlases comprised of tens of thousands of cells primarily focus on the *Arabidopsis thaliana* (herein referred to as Arabidopsis) root, and they provide refined insight to well-characterized cell type-specific lineage progressions (Denyer et al., 2019; Ryu et al., 2019; Shahan et al., 2020; Shulse et al., 2019; Zhang et al., 2019). While these pioneering studies have validated scRNA-seq approaches for resolving unidirectional differentiation processes, they have not focused on flexible and indeterminate features of plant development. One exemplary developmental model for cell fate specification and flexible lineage progression is the Arabidopsis epidermal stomatal lineage (Pillitteri and Torii, 2012). Stomata are cellular valves consisting of two sister “guard cells” that modulate the aperture of a pore between them to facilitate atmosphere gas exchange. Guard cells are one of the final products of a multipotent stem cell lineage that gives rise to the leaf epidermis. Initial stages of the lineage feature asynchronous and indeterminate divisions modulated by extrinsic environmental cues, enabling cell fate flexibility to generate the requisite density and pattern of stomata for a given environment (Hetherington and Woodward, 2003). Given that environmental cues integrate with underlying genetic programs, it is thought that core regulators serve dynamic roles within cell states to tune cell specification and differentiation (Lau et al., 2018; Vatén et al., 2018), though it is unclear how the flexibility of cell fates is controlled.

Here, we leveraged scRNA-seq and molecular genetics to examine dynamic developmental states of the stomatal lineage in the context of the growing Arabidopsis leaf. Examining stomatal production and lineage progression within a broader organ-level context provides insight into cellular programs that coordinate to build a functional organ. scRNA-seq identified distinct models of cell state differentiation within leaf tissue, revealing cell states along with putative regulators across the mesophyll, vasculature, and epidermis. Our atlas of leaf development serves as a useful resource to pursue creative questions and generate hypotheses, as well as inspire comparative analyses with other organs or species, with the goal to better define attributes of tissue development both within an organism and between species. Applied to the question of developmental flexibility, scRNA-seq revealed that flexible cell states on the epidermal landscape may exist along a continuum, with progressive cell specification. This model suggests that obligate core transcriptional regulators and signaling cascades control a range of cellular programs within lineage continuums. One such critical regulator, the transcription factor SPEECHLESS (SPCH), is known to integrate environmental fluctuations in the early stages of the lineage. However, our scRNA-seq and subsequent functional perturbations indicated that SPCH serves a broadened role that enforces cell specification and commitment to generate stomata. Collectively, our findings uncover progressive cell state decisions along lineage trajectories in developing leaves.

## RESULTS

### Complementary scRNA-seq approaches to resolve underlying cell heterogeneity

To leverage our understanding of flexible divisions within the epidermal stomatal lineage, while also surveying developmental processes within inner tissue mesophyll and vasculature, we used complementary single-cell RNA sequencing (scRNA-seq) approaches (Table S1) with 10 days-post-germination (dpg) seedlings. Overall, we sequenced transcriptomes from ∼18,000 cells with droplet-based 10X Genomics and ∼500 cells with plate-based Smart-seq2. This approach allowed us to capture a range of cell states, gene coverage, and cytometry data. In all cases, prior to sequencing, whole leaf tissue was broken down by enzyme-mediated protoplasting followed by fluorescence-activated cell sorting (FACS), which enriched for rare stomatal lineage cells that are challenging to capture from whole tissue. We ultimately resolved underlying heterogeneity among cell states in the developing leaf. Here, we define cell states as instances of putative cell transitions or types (e.g. specific precursor or mature cell) and thereby encompass the range of possibilities for what it means to be a cell along a lineage trajectory (Morris, 2019). Insight into cell states enables us to build models of differentiation programs delineated by predicted lineage trajectories.

### Distinct models of cell state differentiation within leaf tissue

We first profiled cell states from seedlings that expressed the transgene *ML1p::YFP-RCI2A*, in which YFP is fused to the transmembrane domain-containing protein RCI2A and expressed under the predominantly epidermal homeobox transcription factor *MERISTEM LAYER 1 (ML1)* (Lu et al., 1996; Nylander et al., 2001; Roeder et al., 2010). It should be noted that the *ML1* gene promoter is also lowly active in internal tissue cells (Iida et al., 2019), and previous studies showed that FACS-enrichment with this promoter isolates mesophyll cells among other cell types (Adrian et al., 2015; Tian et al., 2019; Yadav et al., 2014). Thus, while our approach enriched epidermal cells at the expense of the more abundant mesophyll cells, the sorted pool of cells was expected to contain a mix of epidermal and inner-tissue cells that would enable comparative analyses. Single cells were captured by droplet-based 10X Genomics microfluidics (Zheng et al., 2017) for downstream library preparation and sequencing. We built our gene expression matrix with Cell Ranger and used the Seurat v3 analysis pipeline (Butler et al., 2018) to obtain transcriptomes for 5,021 cells, with a median of 1,870 detected genes and 5,026 unique molecular identifiers (UMIs) per cell (Figure S1). Our scRNA-seq atlas of the developing leaf offers a substantial number of detected genes and cell expression profiles to uncover putative developmental programs.

Graph-based unsupervised clustering and differential expression analysis revealed potential cell states within mesophyll, vasculature, and epidermal tissues (Figure 1A, Table S5). Clusters of cells were visualized on a Uniform Manifold Approximation and Projection (UMAP) plot, wherein local and global graph structure is preserved to represent the similarity of cell transcriptomes within and between clusters (Becht et al., 2018; McInnes et al., 2018). While the analysis can be performed at distinct resolutions to yield broader or refined clusters (Figure S2, Figure 1A), we primarily focused on a refined resolution to enable examination of putative cell states. At the developmental stage profiled, there are proliferating cells in all aerial tissues of the seedling. The transcriptional signature of cell cycle phases can override and confound cell state assignment (Butler et al., 2018), so we performed a regression to correct for this and enable comparisons of related modules within different tissues (Table S4, Figure S1).

**Figure 1.**
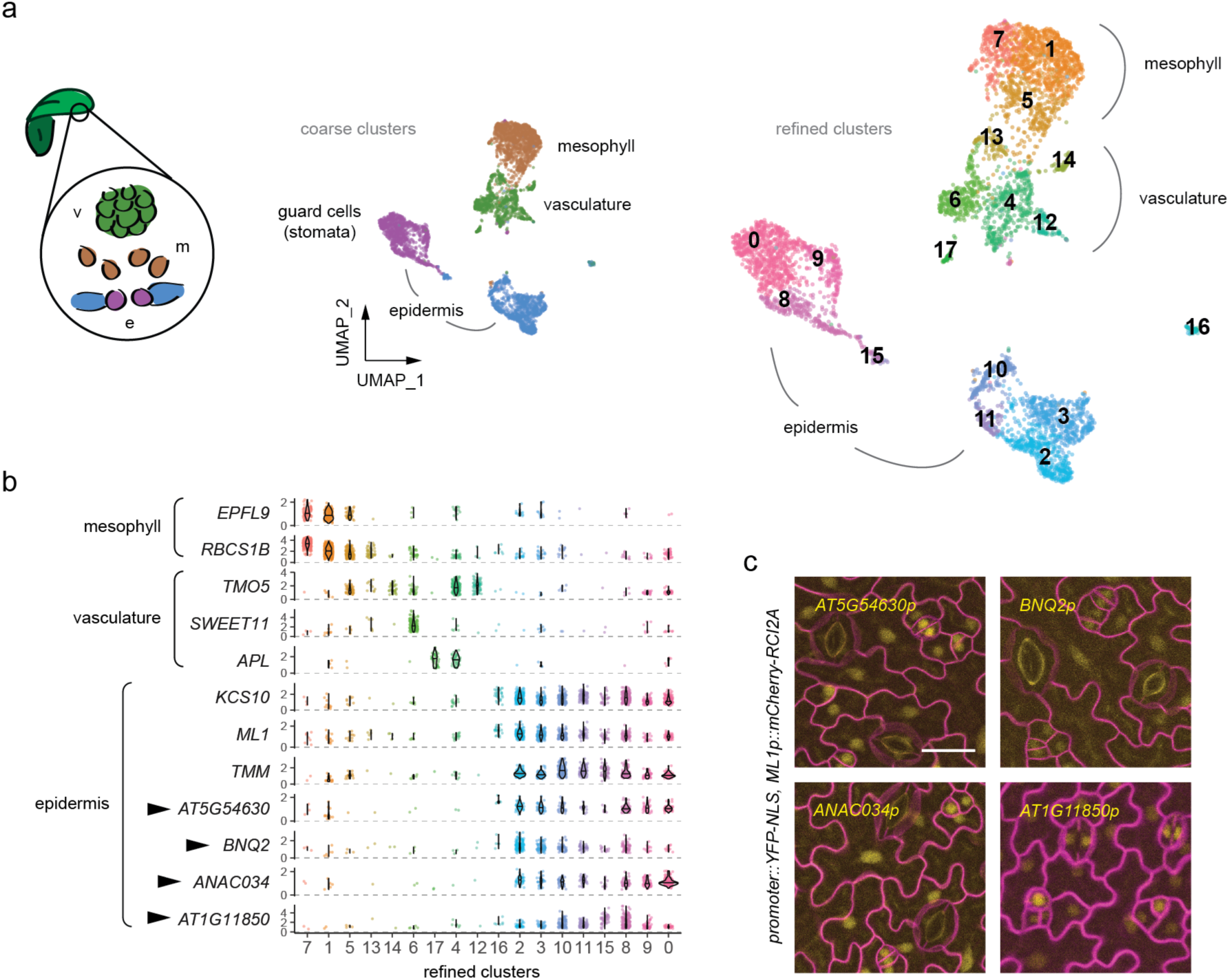
scRNA-seq atlas of the developing leaf. (a) Schematic of a leaf cross-section (left, m: mesophyll, v: vasculature, e: epidermis) is color-coded to match the UMAP of developing leaf tissue from scRNA-seq (10X Genomics v2, 5,021 cells displayed) with 10 days-post-germination (dpg) *ML1p::YFP-RCI2A* seedlings. With a coarse resolution (left, resolution 0.1), major clusters from the mesophyll, vasculature, and epidermis tissue layers are distinguishable. Fine resolution clusters (right, resolution 1) of putative cell states within our UMAP atlas underscore the possibility of capturing state transitions within lineages. Colors and respective number labels represent distinct clusters from graph-based clustering, and number labels designate the order from the largest to smallest cluster (0 to 17, respectively), based on the number of cells per cluster. (b) Violin plots depict expression profiles of known and predicted tissue-specific players within resolution 1 clusters, colored accordingly. Arrowheads correspond to expression profiles of genes with unappreciated roles in the epidermis, for which reporters were generated in (c). (c) Confocal images of promoter-driven *YFP-NLS* reporters (yellow) corroborate predicted profiles from our developing leaf UMAP atlas. Cell outlines (magenta) are visualized with *ML1::mCherry-RCI2A*, and the abaxial epidermis was imaged from first true leaves of 10 dpg seedlings. All images were taken at the same magnification (scale bar: 20 µm).

We began by broadly defining coarse cell clusters that correspond to mesophyll, vasculature, and epidermal cells, based on the expression of well-characterized, tissue-specific players (Figure 1A-B). Similar to findings in Arabidopsis roots (Shulse et al., 2019), many genes with known roles in leaf development were enriched in our differential expression analysis, but were not necessarily the most enriched genes (Table S5). Mesophyll cells were demarcated by expression of the secretory peptide gene *STOMAGEN* (also known as *EPIDERMAL PATTERNING FACTOR LIKE 9, EPFL9*) (Hunt et al., 2010; Kondo et al., 2010; Sugano et al., 2010) and the photosynthetic enzyme RuBisCO subunit gene *RBCS1B*, among other genes encoding for proteins involved in photosynthesis (Sawchuk et al., 2008). Vasculature cells were demarcated by genes encoding regulators of the xylem and phloem, respective water and sugar transporting tissues. These included the xylem bHLH transcriptional regulator gene *TARGET OF MONOPTEROS 5 (TMO5)* (De Rybel et al., 2013), along with the phloem sugar transporter gene *SWEET11* (Chen et al., 2012) and MYB transcriptional regulator gene *ALTERED PHLOEM DEVELOPMENT (APL)* (Bonke et al., 2003). Epidermal cells were demarcated by broad expression of the homeobox transcriptional regulator gene *ML1*, along with genes encoding the cuticle biosynthesis enzyme *3-KETOACYL-COA SYNTHASE 10 (KCS9/FDH)* (Pruitt et al., 2000; Yephremov et al., 1999) and the stomatal lineage leucine-rich repeat (LRR) receptor-like *TOO MANY MOUTHS (TMM)* (Nadeau and Sack, 2002). We found that mesophyll and vasculature cell identities lie adjacent to each other within our developing leaf atlas, while younger epidermal cells are separated from differentiated guard cells. To further validate our atlas, we generated in planta *YFP-NLS* reporters with promoters from a selection of differentially expressed genes (Figure 1B-C, Figure S3), which provided additional support for our epidermal cluster assignments.

To elucidate potential cell states within the mesophyll, vasculature, and epidermis, we inferred transcriptional dynamics with a steady-state deterministic model using scVelo (Figure 2A) (Bergen et al., 2019; La Manno et al., 2018). Briefly, the approach predicts future cell states by leveraging ratios of nascent (unspliced) and mature (spliced) RNA transcripts. In our sequencing data, we detected reads of unspliced transcripts from secondary priming events during 10X Genomics library construction (5% of total reads: ∼3,500 unspliced reads/cell with ∼70,000 total reads per cell). We therefore were able to identify putative, directed transitions between cell states in our atlas, which were corroborated by previously described expressed genes (Figure 2B-E), even though the scVelo model relies on rarely detected transcripts and cell states. We also identified enriched components of signaling pathways and transcriptional control that define these transitions and states in the mesophyll, vasculature, and epidermis (Table S5). Our atlas thus has the potential to yield insight into developmental lineage trajectories that have eluded previous analyses.

**Figure 2.**
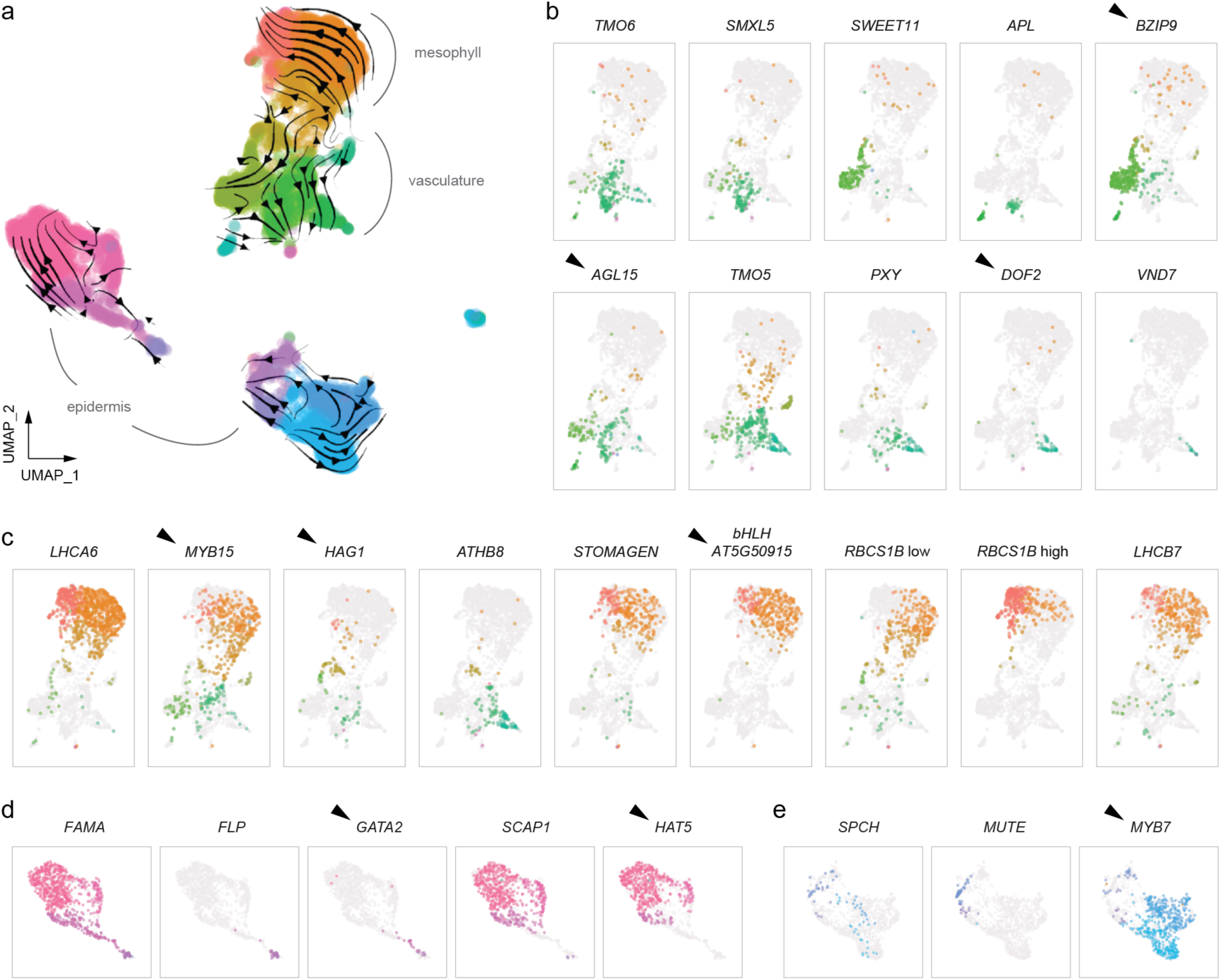
Models of cell state differentiation within leaf tissue. (a) Lineage inference models by scVelo layered onto the UMAP of the developing leaf scRNA-seq atlas. Size of arrows indicate the degree of gene expression changes within predicted, directed transitions from one cell state to another. Colors represent graph-based clustering with resolution 1, and grey brackets demarcate major clusters from the mesophyll, vasculature, and epidermis. (b-e) UMAP insets of the major clusters that correspond to different tissues; which include (b) vasculature, (c) mesophyll, (d) stomatal guard cells, and (e) young epidermal cells. Expression of respective dynamic lineage genes is highlighted by resolution 1 cluster colors. Two insets depicting *RBCS1B* expression distinguish ‘low’ versus ‘high’ gene expression within mesophyll cells. Cells with ‘low’ and ‘high’ expression were selected from the bottom and top 25%, respectively. Representative, previously uncharacterized, potential regulators are designated by arrowheads.

**Figure 3.**
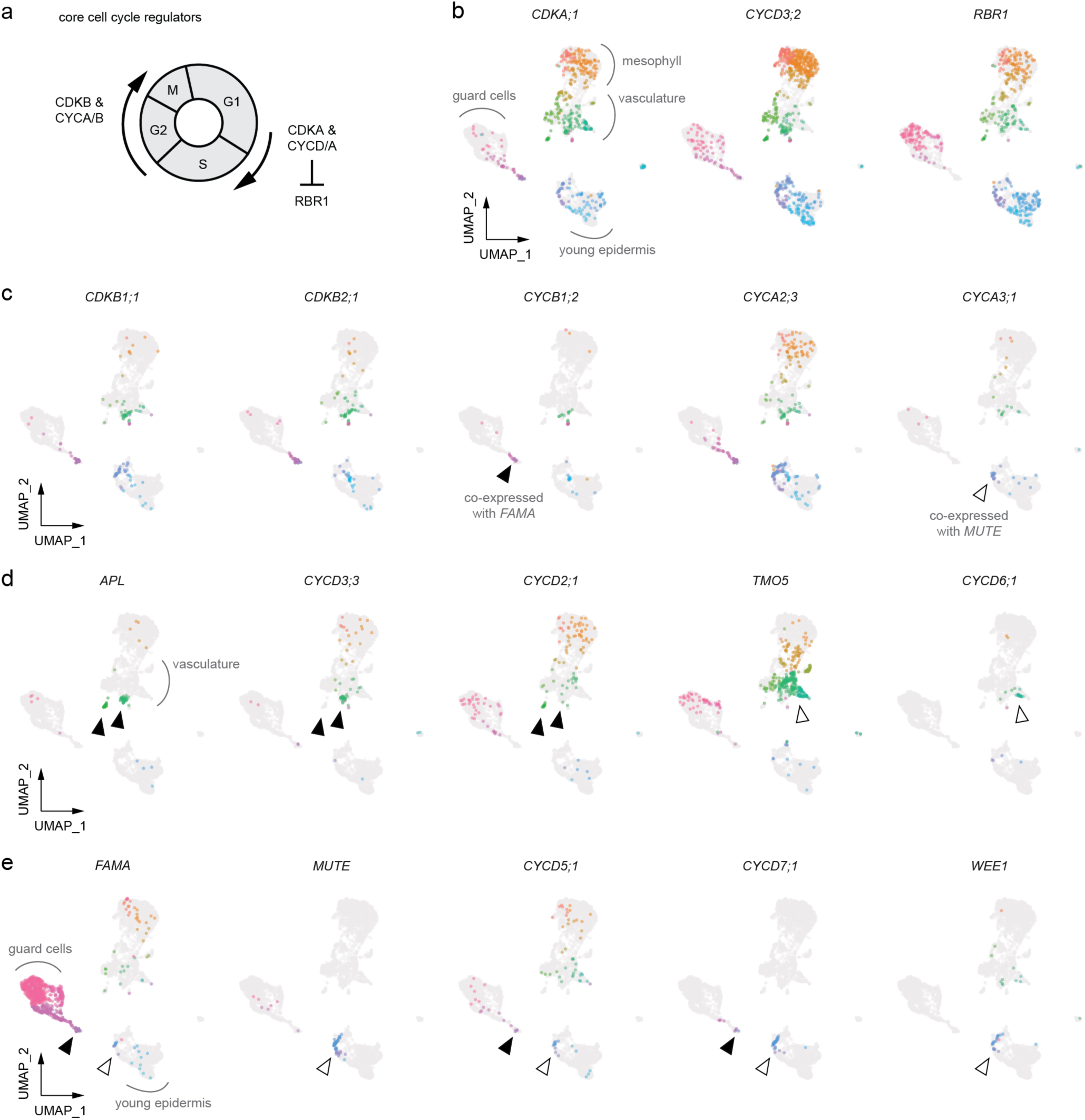
Cell cycle regulators exhibit distinct, yet overlapping expression profiles. (a) Schematic of the cell cycle highlights the regulatory roles of cyclin-dependent kinases (CDKs) and cyclins (CYCs), as well as the G1/S-phase checkpoint regulator RETINOBLASTOMA-RELATED PROTEIN 1 (RBR1). (b-e) UMAPs depict gene expression profiles of cell cycle regulators within the developing leaf atlas. Colors represent graph-based clustering with resolution 1. (b) Core regulators of G1/S phase. (c) *CDKB/CYCB* markers of G2/M phase, along with distinct expression profiles of *CYCAs*. While *CYCA2;3* expression is relatively broad, *CYCA3;1* is likely expressed only in G1/S phase, along with the epidermal transcriptional regulator *MUTE*. (d) Co-expression of *CYCD* regulators with the phloem transcriptional regulator *APL* (black arrowheads) and the procambium/xylem transcriptional regulator *TMO5* (white arrowhead). (e) Co-expression of *CYCDs* and *WEE1* with the epidermal transcription factors *FAMA* (black arrowhead) and *MUTE* (white arrowhead).

Genetic programs that drive specification and differentiation of vascular xylem and phloem tissues are well defined in root, stem, and hypocotyl tissue (Hellmann et al., 2018), and thus provide a useful framework to evaluate the scVelo lineage inference trajectory. Using known markers, we identified procambial meristematic cells, vasculature precursor cells, and maturing phloem and xylem cells (Figure 2B). The developing leaf vasculature exhibited a similar trajectory as in other tissues; expression of the transcriptional regulator genes *SUPPRESSOR OF MAX2-LIKE* 5 (*SMXL5*) and *TARGET OF MONOPTEROS 6* (*TMO6/DOF5*.*3*) (Wallner et al., 2017) demarcated differentiating procambium and phloem, while the sugar transporter gene *SWEET11* (Chen et al., 2012) defined potential companion and parenchyma cells, and the MYB family transcriptional regulator gene *APL* was expressed in mature phloem cells, downstream of the predicted onset of *SMXL5* and *SWEET11*. The bHLH transcriptional regulator gene *TMO5* was expressed in the procambium and overlapped with *SMXL5* expression in our atlas. It was also expressed in the xylem, along with the receptor-like kinase gene *PHLOEM INTERCALATED WITH XYLEM (PXY)* (Fisher and Turner, 2007). Downstream of *TMO5* and *PXY* expression in our lineage inference, the transcriptional regulator gene *VASCULAR RELATED NAC-DOMAIN PROTEIN 7 (VND7)* (Kubo et al., 2005) was restricted to differentiating xylem cells with activated secondary wall biosynthesis programs. Our differential expression analysis identified several putative transcription factors with unappreciated roles in the vasculature (e.g. *BASIC LEUCINE ZIPPER 9, BZIP9; AGAMOUS-LIKE 15, AGL15; and DOF ZINC FINGER PROTEIN 2, DOF2)* (Table S5). A comparison with root scRNA-seq data (Ryu et al., 2019) indicated similar restriction of expression to phloem or xylem cell types. Further analysis is necessary to define the function of these genes, though as a general strategy for prioritizing potential lineage regulators, there is weight in the consistent association of a gene with common cell types from different tissues. Conversely, identification of regulators specific to leaf vasculature cells, compared with the root and other tissues, may point to respective divergent developmental processes.

Given that we were able to elucidate vasculature cell states, we next asked whether our developing leaf atlas could provide insight into mesophyll differentiation. Mesophyll function in photosynthesis is extensively studied, but less is known about cell differentiation states in this considerably less accessible and tractable inner leaf tissue. Trajectory inference indicated that distinct cell fate decisions distinguish the mesophyll and vasculature (Figure 2C). Clusters of cells from the mesophyll and vasculature appear bridged by putative ground meristem and procambium cells that give rise to differentiating cells within each tissue. For instance, onset of the light-harvesting complex gene *LHCA6* and the homeobox transcription factor gene *ATHB8*, previously characterized progenitor markers (Sawchuk et al., 2008; Scarpella et al., 2004), was detected in ground cells that are committed to mesophyll and vasculature development, respectively. We thus leveraged differentially expressed genes (Table S5) to identify enriched putative regulators of the ground meristem, which yielded the transcription factor genes *MYB15* and *HAG1*/*MYB28*. Conversely, analysis of putative mesophyll differentiation regulators identified the bHLH transcription factor gene *AT5G50915*, that also overlapped with the secretory peptide gene *STOMAGEN/EPFL9* involved with early epidermal-mesophyll tissue coordination to control stomatal density (Hunt et al., 2010; Kondo et al., 2010; Sugano et al., 2010). Interestingly, we may be able to distinguish the adaxial palisade and abaxial spongy mesophyll, based on respective divergent expression of the light-harvesting complex gene *LHCB7* and the RuBisCO subunit gene *RBCS1B* (Sawchuk et al., 2008), though there is substantial overlap between these expression profiles. One feature of *RBCS1B* expression, as with other highly expressed genes, was that it also enabled us to distinguish a quantitative gradient within our scRNA-seq data, with low expression detected in cells predicted to be precursors of those with higher expression. Through new perspectives on establishment and maintenance of cell states, our atlas may extend our understanding of palisade and spongy mesophyll differentiation.

Lineage inference also provided insight into directed developmental transitions within the epidermis. Epidermal cells were divided into two main clusters in our atlas. One cluster featured expression of the bHLH transcriptional regulator gene *FAMA*, which indicated that it represents differentiating stomatal guard cells (Figure 2D). Within this cluster, the MYB transcription factor gene *FOUR LIPS (FLP/MYB124)* exhibited much more restricted expression, as previous reports link *FLP* to the earliest stages of guard cell differentiation (Lai et al., 2005). Restricted to the opposite region within the cluster was the Dof-type transcriptional regulator gene *STOMATAL CARPENTER 1 (SCAP1)* (Negi et al., 2013), indicating that these are mature guard cells. With *FAMA, FLP*, and *SCAP1* gene profiles as guides of differentiation states, differential gene expression analysis identified additional putative regulators, such as the transcription factors *GATA2* and *HAT5* that respectively mark the beginning and end of guard cell differentiation.

The second epidermal cluster in our atlas represented young epidermal and stomatal lineage cells (Figure 2E). A notable feature of this cluster was its diverging trajectory inference (depicted as deviating arrows on the UMAP, Figure 2A) modeling multiple cell fate decisions. A finer examination of these states that includes additional scRNA-seq data follows below (Figure 4B), but at this resolution, we observed relatively broad expression of the bHLH transcriptional regulator gene *SPCH* (MacAlister et al., 2007; Pillitteri et al., 2007) in early stomatal lineage stem cells. Its paralogue *MUTE* (MacAlister et al., 2007; Pillitteri et al., 2007) displayed much more restricted expression in cells committing to stomatal guard cell differentiation, and accordingly, these cells in the young epidermal cluster were oriented toward the guard cell cluster. Opposed to *MUTE*, cells expressed the transcription factor gene *MYB7*. MYB7 has been associated with production of secondary metabolite phenylpropanoids (Albert, 2015) and may define cells progressing toward the alternative pavement cell fate. Genes encoding cuticular wax biosynthesis and defense-related factors were co-expressed with *MYB7* (Table S5), and overall there were a substantial number of putative regulators dynamically expressed in patterns traversing the young epidermal clusters. Thus, in contrast with previous RNA-seq studies that were biased by bulk identification of cells expressing specific markers (e.g. *SPCH, MUTE*, and *FAMA*) (Adrian et al., 2015), leveraging the broadly expressed *ML1* gene promoter with scRNA-seq enabled us to capture cells that take alternative trajectories to generate either stomata or pavement cells. We thus resolved heterogeneous transcriptional changes and distinct models of cell state differentiation on the epidermis of developing leaves.

**Figure 4.**
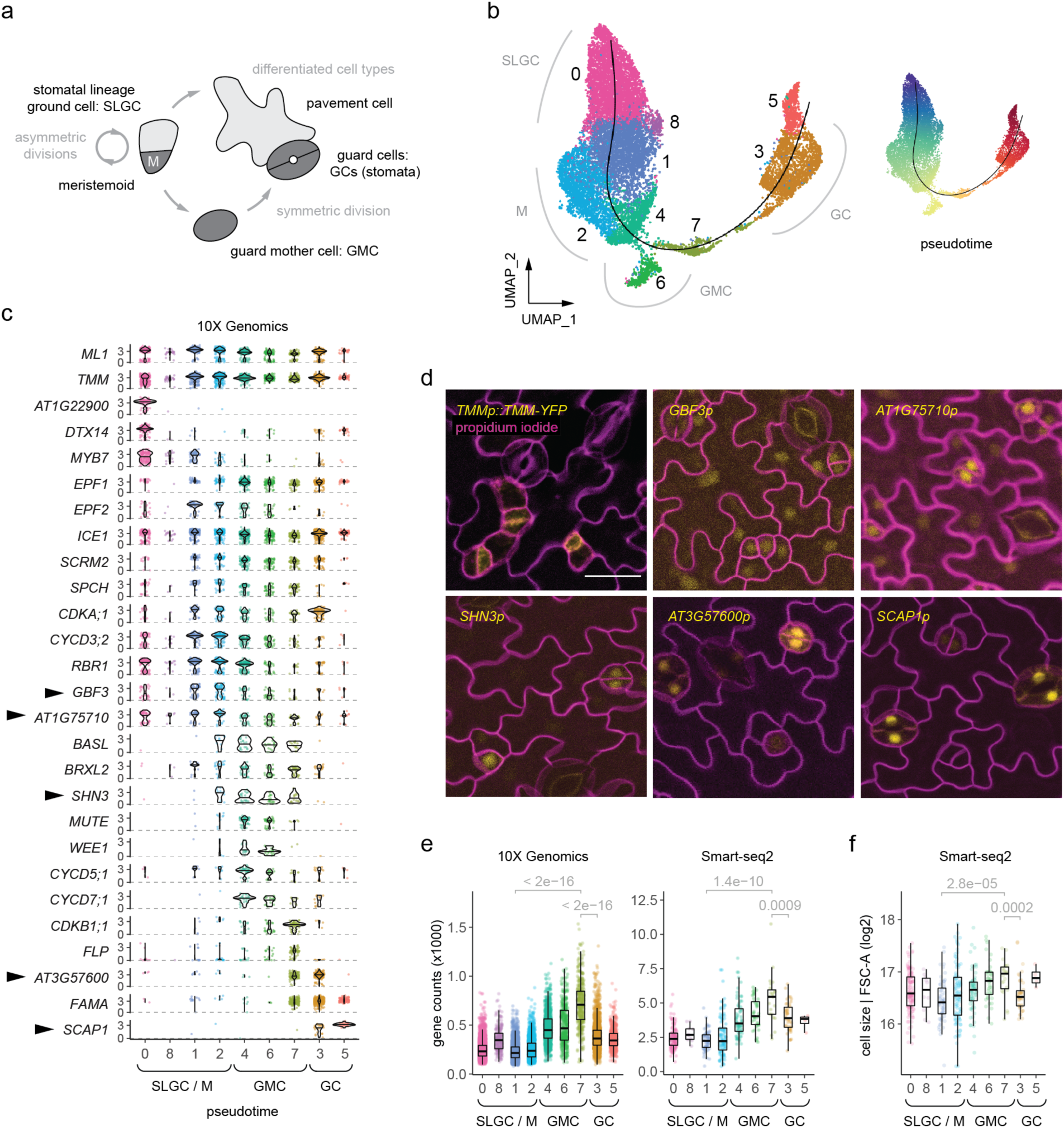
Stomatal lineage flexibility defined as a continuum of cell states. (a) Model of stomatal lineage highlights that early stem cells, meristemoids (Ms) and stomatal lineage ground cells (SLGCs), either divide asymmetrically (left) or commit to differentiation (right). Differentiation programs ultimately yield guard cells (GCs) and pavement cells. (b) UMAPs of integrated scRNA-seq (10X Genomics v3, 12,933 cells; and Smart-seq2, 478 cells) derived from *TMMp::TMM-YFP*-expressing shoot tissue from 10 dpg seedlings. Colors depict graph-based clustering with resolution 0.4 (left) and the relative pseudotime scale (right). Overlaid on each UMAP is the pseudotime axis (black line). (c) Violin plots depict a range of gene expression profiles within pseudotime, aligned with cluster resolution 0.4 (colors). Included are known regulators of the lineage, as well as genes for which reporters were generated (arrowheads). Pseudotime is predicted to initiate with *SPCH* expression, and it represents cell state transitions from Ms to SLGCs in one direction (towards left), as well as transitions from Ms to GMCs and GCs (towards right). (d) Confocal images of promoter-driven *YFP-NLS* reporters corroborate predicted expression profiles. The abaxial epidermis was imaged from first true leaves of 10 dpg seedlings. All images were taken at the same magnification (scale bar: 20 µm). (e) Boxplots depict transcriptional diversity within pseudotime, based on total gene counts from respective datasets. (f) Boxplots depict relative cell size (log2 values of the median forward scatter area, FSC-A) within pseudotime. FSC-A was measured and indexed by FACS, corresponding to the Smart-seq2 dataset. Colors in (e, f) represent cluster resolution 0.4, and clusters are aligned with pseudotime, as in (c). Adjusted p-values in (e, f) are from pairwise comparisons using the Wilcoxon rank sum test.

An emergent signature from our atlas may mark the spatial origin of cells. All tissues in the leaf display adaxial (top) and abaxial (bottom) polarity (Kidner and Timmermans, 2010), which can be seen with the vasculature xylem and phloem, palisade and spongy mesophyll, and the adaxial and abaxial epidermis. Overlaying expression of the core regulators, including members of the *HD-ZIP III, KANADI* and *YABBY* gene families (Figure S4), enabled us to detect expected patterns of the leaf polarity axis in internal tissues. We also found that we have potentially obtained epidermal cells from each side of the leaf. Because expression of such polarity genes is highest in leaf primordia (Ram et al., 2020), we lose the ability to more confidently assign location in maturing leaf cells. Nonetheless, we found a substantial number of guard cells expressing adaxial or abaxial signatures (Table S6), which may provide insight into physiological processes prioritized by stomata originating from these different surfaces of the leaf (Mott, 2007).

### Cell cycle regulators exhibit distinct, yet overlapping expression profiles

We next asked whether comparisons between leaf tissues can serve to define distinct and overlapping features of common regulatory modules. We hypothesized that common modules integrate with and are controlled by different tissue-specific contexts. One such critical, conserved module is the cell cycle. Because the expression signature for G1/S or G2/M phases can overwhelm other features of cell identity, we originally regressed out the cell cycle signature from our analysis (Figure S1). However, plant genomes often encode several core cyclin-dependent kinase (CDK)/cyclin regulators, which can exhibit lineage-specific expression and function (Adrian et al., 2015; Gutierrez, 2016; Han et al., 2018; Sozzani et al., 2010; Weimer et al., 2018). An active area of research aims to understand how cell cycle and differentiation processes are coordinated.

In the leaf atlas, we found that core regulators exhibit distinct, yet overlapping, expression within vasculature, mesophyll, and epidermal cell states (Figure 3A, Figure S5). G1/S-phase transition genes *CYCLIN-DEPENDENT KINASE A;1 (CDKA;1), CYCLIN D3;2 (CYCD3;2), and RETINOBLASTOMA-RELATED PROTEIN 1 (RBR1)* were broadly expressed (Figure 3B), while members of the G2/M-phase *CDKB* and *CYCB* gene families were expressed in distinct clusters (Figure 3C). Of note, we found that *CYCD* gene family members exhibited both broad and tissue-specific expression profiles (Figure 3D-E). While the gene *CYCD3;2* was broadly expressed, *CYCD3;3* and *CYCD2;1* were selectively enriched in distinct mature phloem clusters that expressed the transcriptional regulator *APL*. Conversely, *CYCD6;1* was co-expressed with the procambium/xylem bHLH transcriptional regulator *TMO5*. Onset of *CYCD7;1* in the epidermis exclusively overlapped with the transcriptional regulator *MUTE* and extended into a domain shared with the emergence of *FAMA* expression. *CYCD5;1* was broadly detected in a few cells, and may overlap with *CYCD6;1* and *CYCD7;1* in the vasculature and epidermis, respectively. Interestingly, the kinase *WEE1* is expressed in the epidermal and vascular clusters where restricted *CYCDs* converge (Figure 3E). WEE1 is a negative regulator of the G2/M transition and ensures size control in yeast (Nurse and Thuriaux, 1980). In Arabidopsis, WEE1’s role has been described only in response to DNA damage (Cools et al., 2011; De Schutter et al., 2007; Sorrell et al., 2002). *CYCA* gene family members also displayed a range of expression profiles within the atlas (Figure 3B). For instance, while *CYCA2;3* was broadly expressed, the G1/S-phase *CYCA3;1* (Takahashi et al., 2010) was selectively co-expressed with *MUTE*. A closer examination of cell cycle genes revealed that MUTE may coordinate with G1/S-phase regulators, while FAMA coordinates with G2/M-phase regulators. (Figure 3C, 3E). This is consistent with proposed roles of these transcription factors in driving cell cycle progression (Han et al., 2018; Ohashi-Ito and Bergmann, 2006), though our findings also raise the possibility that *MUTE* and *FAMA* gene expression is reciprocally responsive to cell cycle state.

### Integrated analysis elucidates flexible stomatal lineage cell states along a continuum

As distinct models of cell state differentiation were observed in our scRNA-seq leaf atlas (Figure 2), we next aimed to further elucidate genetic programs within the well-characterized and tractable stomatal lineage. We performed scRNA-seq on FACS-isolated cells expressing the leucine-rich repeat (LRR) receptor-like gene *TOO MANY MOUTHS (TMM; TMMp::TMM-YFP*) (Nadeau and Sack, 2002). TMM is highly expressed in stem cells of the stomatal lineage, including both meristemoids and stomatal lineage ground cells (SLGCs). Such cells are defined as flexible given that they are able to either asymmetrically divide to yield another meristemoid/SLGC pair or differentiate into distinct epidermal cell types (Nadeau and Sack, 2002) (Figure 4A). SLGCs eventually undergo endoreduplication and differentiate into pavement cells, though this mature cell type is not captured in our datasets here due to cell size selection. Conversely, meristemoids differentiate into guard mother cells (GMCs), which divide symmetrically to yield a pair of stomatal guard cells (GCs). While decisions to divide or differentiate are known to be regulated by cell-cell signaling cascades, cell polarity, and transcriptional control (Lee and Bergmann, 2019), previous studies focused on discrete major fate transitions. The extent of cell state heterogeneity across flexible cell identities of meristemoids and SLGCs remains unclear.

We leveraged an integrated scRNA-seq approach focused solely on stomatal lineage cells to address characteristic and generalizable developmental features displayed by the stomatal lineage. These included a potential continuum of early lineage cell states and possible relationships among cell size, transcriptional diversity, and cell commitment during GMC fate commitment. To take advantage of a variety of computational analyses, some that require many cells and others that require deep sequence coverage, we used two different technologies to profile *TMMp::TMM-YFP* expressing young stomatal lineage cells. Cells captured for droplet-based 10X Genomics (Zheng et al., 2017) yielded transcriptomes for 12,933 cells, with a median of 353 detected UMIs and 271 genes. This dataset was integrated with another we generated by plate-based Smart-seq2 (Picelli et al., 2014), which yielded transcriptomes for 478 cells, with a median of 2,786 detected genes. After integration with Seurat v3 (Butler et al., 2018), potential cell states were identified through graph-based clustering and differential gene expression analysis (Figure S6-S7, Figure 4B, Table S7). To identify a putative lineage trajectory model that reflects transitions between cell states, we reconstructed pseudotime with simultaneous principle curves using slingshot (Street et al., 2018). Compared with other methods, slingshot offers relatively more stringent predictions of putative transitions and has been previously identified as a high-performing program (Saelens et al., 2019). Depicted as a smooth curve overlaid on the UMAP, the pseudotime axis revealed potential developmental relationships between cell states.

Cell states were annotated using a similar approach as with the leaf atlas, assessing expression profiles of genes that encode known players in the lineage, along with genes that display dynamic profiles across the clusters (Figure 4C). Flexible cell states early in the lineage (meristemoids and SLGCs) indeed appeared to exist along a continuum within pseudotime, defined by early expression of *SPCH* and *INDUCER OF CBF EXPRESSION1 (ICE1)/SCREAM (SCRM)* bHLH transcriptional regulators (Kanaoka et al., 2008). Separate and distinct clusters represented transitions through deterministic guard cell fate commitment, with the expression of *MUTE* followed by *FAMA*. Surprisingly, *SPCH* expression extended into the expression domains of *MUTE* and *FAMA*, which was unexpected given functional genetic analyses placing *SPCH* early in stomatal lineage progression (Davies and Bergmann, 2014; MacAlister et al., 2007; Pillitteri et al., 2007). We further validated our lineage model of cell state dynamics in vivo. We created reporters of genes with no known function in the lineage, that displayed both broader and specific expression within pseudotime (Figure 4D, Figure S8). Imaging these reporters in 10 dpg leaves corroborated expression profiles from the scRNA-seq lineage trajectory, which indicates that our pseudotime model of cell states may reflect in vivo dynamics.

### Transcript diversity correlates with cell size measured by FACS

We next asked whether cell states within our lineage pseudotime could be leveraged to uncover mechanisms for GMC fate commitment, a deterministic directed process that promotes one symmetric division event to yield a pair of stomatal guard cells. We found that the continuum of early lineage cell states included the onset of GMC commitment (Figure 4B-C). This indicates that there may be a gradual, stochastic process by which meristemoids differentiate. We were thus especially intrigued to discover increased transcriptional diversity within differentiating GMCs (Figure 4E), as this feature of lineage progression has been recently described in a range of animal and human systems (Gulati et al., 2020). Furthermore, stomatal lineage cells range in size, and it remains unclear how this is regulated. We therefore examined whether increased transcriptional diversity was correlated with cell size. Using FACS forward scatter data with indexed cells captured in our Smart-seq2 scRNA-seq analysis, cell size was positively correlated with transcriptional diversity; it gradually increased with the onset of GMC fate commitment in pseudotime, with a drop-off in size after the inferred symmetric division (Figure 4F). This is in contrast to the early lineage clusters where a range of cell sizes were detected without a clear correlation with transcriptional diversity, as expected for flexible cell states among asymmetrically dividing cells.

Given the observation of increased transcriptional diversity, we asked whether specific transcription factors and chromatin remodelers were expressed during the onset of GMC fate commitment to modulate the transcriptional landscape. As seen in here (Figure 4C, 4F), *MUTE* expression is detected in GMCs of all sizes, but *FAMA* is limited to only the largest GMCs. In addition to activators iteratively expressed to gradually activate downstream programs, we might expect a requirement for negative regulators of the lineage progression repressors. We therefore identified putative transcriptional regulators that were differentially expressed within pseudotime (Table S7). Our stomatal lineage scRNA-seq dataset thereby reveals potential candidates that will serve as a framework for future studies to examine underlying mechanisms that lock flexible cell states into a given deterministic fate.

### Broadened functional role of an early-lineage core regulator

A paradigm of Arabidopsis stomatal development is that sequential expression of *SPCH, MUTE*, and *FAMA* defines discrete transitions of cell fates (MacAlister et al., 2007; Ohashi-Ito and Bergmann, 2006; Pillitteri et al., 2007). However, there are indications from our lineage pseudotime described here (Figure 4B-C) that developmental progression is more complicated. In our gene expression overlays of *SPCH, MUTE* and *FAMA*, expression of *SPCH* persisted beyond early stages of the lineage (Figure 4C). Distinct cell states even appear to co-express *SPCH* with either *MUTE* or *FAMA* (Figure 5A). Given the potential conflict of these findings with the sequential model of the lineage, we reexamined *SPCH* expression in vivo within dividing GMCs by time-lapse imaging. We imaged a previously published reporter (coding sequence *SPCHp:cdsSPCH-YFP, cdsSPCH*) (Davies and Bergmann, 2014; Vatén et al., 2018), as well as a new reporter that contained more of the SPCH genomic context (genomic sequence *SPCHp:gSPCH-YFP, gSPCH*), which may more accurately recapitulate regulatory control at the endogenous locus. Each reporter was characterized in a *spch-3* (null) background. Expression of the *SPCH* reporters indeed extended beyond early stage, asymmetrically dividing cells (in 4 dpg cotyledons), with stronger signal potentially detected using the *gSPCH* genomic reporter (Figure 5B). This ultimately corroborates our new model based on scRNA-seq and suggests that SPCH serves an unappreciated role within dynamic cell states.

**Figure 5.**
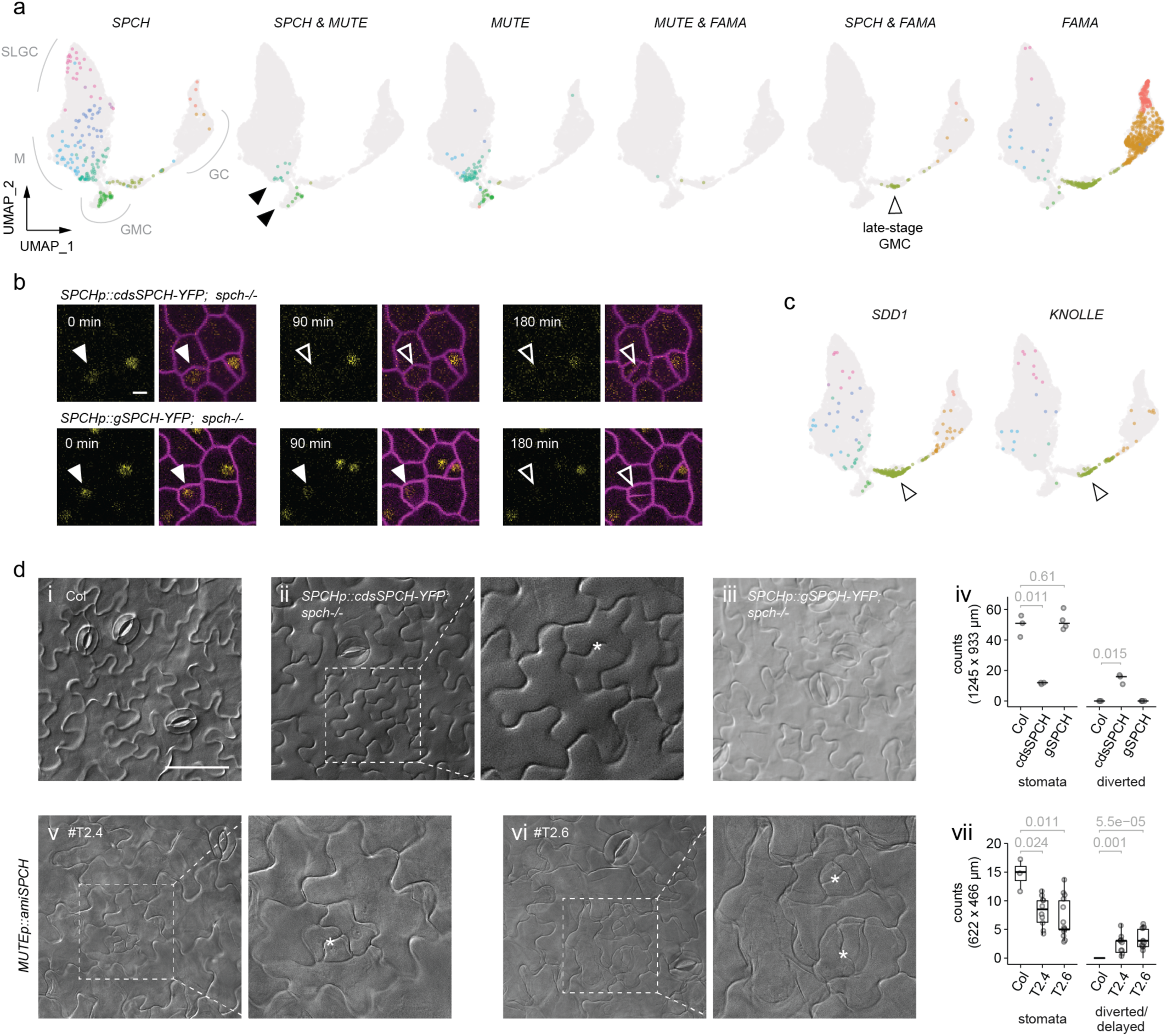
*SPCH* is expressed and required beyond the early stages of the lineage. (a) Extended *SPCH* gene expression is depicted. UMAPs indicate cells and clusters (resolution 0.4) that express combinations of the genes *SPCH, MUTE*, and *FAMA*. Clusters that express both *SPCH* and *MUTE* (black arrowheads) are distinct from the cluster that expresses *SPCH* and *FAMA* (white arrowhead). (b) Time-lapse confocal images of *SPCHp::cdsSPCH-YFP* (protein-coding sequence) and *SPCHp::gSPCH-YFP* (genomic sequence), each complementing *spch-3*. Images are from the abaxial epidermis of 4 dpg seedlings, and arrowheads follow the GMC division that yields young stomatal guard cells (at 180min). Outside the frame of the time-lapse, young guard cells are presumed to enlarge and form a pore. Filled white arrowhead: reporter expression detected, outlined arrowhead: no expression detected. All images were taken at the same magnification (scale bar: 5 µm). (c) Examples of genes, and their respective UMAP expression profiles (colors: cluster resolution 0.4), that may be regulated by SPCH beyond early stages of the lineage. Putative state-specific direct targets were found to be bound by SPCH and differentially expressed within pseudotime. (d) Diverted cell fates (asterisks: borders of diverted/delayed cell identities) in representative differential interference contrast (DIC) images of abaxial cotyledon epidermis from Col wildtype control (i), *SPCHp::cdsSPCH-YFP spch-/*- (ii), *SPCHp::gSPCH-YFP spch-/-* (iii), and two independent lines of *MUTE::amiSPCH* T2.4 (v) and T2.6 (vi). Images were all taken at 15 dpg with the same magnification (scale bars: 100 µm), except for insets demarked by white dashed boxes. Diverted/delayed fate events are quantified in (iv, vii). Diverted fates are defined by presumed exit of the stomatal lineage, wherein sister cells undergo lobing – rather than give rise to a meristemoid and SLGC pair or generate a pair of guard cells. Early lobing of an SLGC is depicted in (vi, top asterisk). *MUTE::amiSPCH* epidermis also displayed delayed lineage commitment, defined by putative additional rounds of asymmetric divisions before the commitment to yield guard cells. Plots in (iv, vii) include grey points that each represent counts from an individual seedling (at least n=3 seedlings were measured per genotype). median: black line. p-values: two-sample t-test for respective comparisons except for diverted events in (iv), wherein the p-value represents one-sample t-test.

To examine the diversity of potential modules regulated by SPCH, we identified putative target genes from ChIP-seq (Lau et al., 2014) and enriched gene ontologies across cell states in the stomatal lineage scRNA-seq dataset. We found that many differentially expressed genes may be directly regulated by SPCH (i.e. bound by SPCH), and we thereby detected similar enriched modules from gene ontology, spanning from stimulus and immune responses to regulation of growth and division processes (Table S7). We also identified genes that correlate with induced gene expression of *SPCH*, by leveraging a previous RNA-seq dataset (Lee et al., 2019). Differentially expressed genes in the clusters that are bound by SPCH and positively correlate with induced *SPCH* expression were generally enriched in early lineage state clusters (Table S7). Of these, within the late-stage GMC cluster, we found that SPCH appears to control genes implicated in cell fate decisions and cell cycle control (Figure 5C), which include the protease gene *STOMATAL DENSITY AND DISTRIBUTION 1* (*SDD1*) (von Groll et al., 2002) and the cytokinesis-specific syntaxin gene *KNOLLE* (Lauber et al., 1997; Lukowitz et al., 1996). Conversely, genes that negatively correlate with *SPCH* expression were detected in the guard cell clusters. As SPCH function likely shifts within different cellular states, it is interesting to consider the potential new role of SPCH in GMCs, where it appears to sequentially overlap with both *MUTE* and *FAMA* expression.

Given that we corroborated extended expression of *SPCH* and identified potential modules putatively controlled by the regulator across cell states, we tested whether extended expression of *SPCH* was required for proper cell identity and fate decisions. The first hint that SPCH might contribute to later differentiation steps came from examination of mature tissue (15 dpg cotyledons) in lines where the *spch-3* null mutant was rescued by *cdsSPCH* or *gSPCH*. While *gSPCH* leaves exhibited similar stomatal counts and broad cellular features compared with wildtype, *cdsSPCH* leaves had fewer stomata, and displayed a novel phenotype, which we call “diverted” stomatal lineage cells based on morphology and position relative to other epidermal cells (Figure 5D, Figure S9). Diverted cells undergo lobing, rather than yield either meristemoid and SLGC pairs or pairs of guard cells. This finding could be due to disrupted levels and/or timing of *SPCH* expression, particularly since the *cdsSPCH* transgene was more lowly expressed at later stages relative to *gSPCH* (Figure 5B). We therefore examined whether SPCH is specifically required at this later developmental transition by selectively depleting late-stage expression with the *MUTE* gene promoter driving artificial microRNAs against *SPCH* (*MUTEp::amiSPCH)*. With these lines, we identified two phenotypes: a delay in commitment (as evidenced by additional rounds of early lineage divisions before producing stomata) and diverted cell identities, similar to what we found in the *cdsSPCH* rescue lines (Figure 5D, Figure S9). Diverted and delayed cellular developmental outcomes were not observed in control seedlings. Overall, this functional approach indicates that fine-tuned expression of *SPCH* is required to control and promote late cell fate commitment in GMCs, and thereby, along with proposed modules from functional genomics, reveals a new role for the critical transcriptional regulator.

## DISCUSSION

Plants display extreme biological flexibility that offers insight into exceptional developmental strategies. Here, we uncovered distinct models of cell state differentiation within Arabidopsis leaf tissue by leveraging single-cell transcriptomics and functional molecular genetics. A comparative analysis of cell states in the mesophyll, vasculature, and epidermal clusters revealed tissue-specific modules and enriched cellular programs, along with adaxial/abaxial tissue polarity signatures. Our scRNA-seq atlas of the developing leaf thereby serves as a useful resource to pursue questions and generate hypotheses that will unravel how these tissues arise. We also identified underlying heterogeneity within cell states of the flexible epidermal stomatal lineage, which appear to exist along a continuum, with progressive cell specification. Our findings indicated that the core early transcriptional regulator SPCH potentially controls a range of cellular programs within the lineage continuum. Early roles may be foundational to adjusting stomatal lineage expansion in line with environmental fluctuations (Kumari et al., 2014; Lau et al., 2018). scRNA-seq analysis showed, unexpectedly, that *SPCH* was expressed beyond the early stages of the lineage, and we demonstrated that it was required for cell fate commitment to yield stomatal guard cells. Overall, our work has refined the stomatal lineage paradigm.

SPCH serves an unappreciated role in promoting stomatal guard cell fate, which is in conflict with the previous model of sequential and discrete control by SPCH, MUTE, and FAMA (MacAlister et al., 2007; Ohashi-Ito and Bergmann, 2006; Pillitteri et al., 2007). When SPCH was specifically depleted by *amiRNAs* driven by the *MUTE* promoter, we detected extended asymmetric amplifying divisions that may indicate a failure to commit and a delay in lineage progression. We also observed that some divisions, rather than resulting in a guard cell pair, yield a pair of lobed cells that are presumed to have diverted from the guard cell fate genetic program. Extended amplifying divisions have also been detected in the *mute* null, though they represent a distinct phenotype, as such divisions are defined by rounded cells within *mute* leaves (MacAlister et al., 2007; Pillitteri et al., 2007). Diverted cell fates as observed in the *MUTEp::amiSPCH* leaves have not been described for other loss-of-function mutations, but ectopic overexpression of the secretory peptide gene *EPIDERMAL PATTERNING FACTOR 1 (EPF1)* leads to a similar phenotype (Hara et al., 2009, 2007; Triviño et al., 2013). Consistent with this, EPF1 represses SPCH function through kinase signaling cascades (Lampard et al., 2008). Thus, in our new model of the stomatal lineage, we postulate that SPCH serves dynamic roles during early flexible cell states, and also complements MUTE and FAMA at later stages to drive cell fate commitment and differentiation of guard cells. Signaling and regulatory modalities that selectively implicate SPCH may be recruited into these later stages. For example, when Arabidopsis is subjected to environmental stresses (e.g. heat, starvation, or drought), the stomatal lineage response has centered on transcriptional and post-translational control of SPCH (Han et al., 2020; Kumari et al., 2014; Lau et al., 2014). The preference for SPCH was often ascribed to its role in early, flexible development. However, SPCH protein also has unique domains that render it a partner or target of regulatory proteins that MUTE and FAMA cannot engage (Lampard et al., 2008; Putarjunan et al., 2019), and thus it may be that SPCH is necessary to transmit such environmental information to later decisions.

Our work also highlighted distinct and overlapping features of common regulatory modules within a developing leaf. We found that cell cycle regulators integrated with tissue-specific regulatory contexts and may coordinate cell fate. For example, we see parallels to the stomatal lineage in the vasculature cluster, two places where cell states are dependent on complexes containing RBR1 and a lineage-specific transcription factor (FAMA and SCARECROW (SCR), respectively) (Cruz-Ramírez et al., 2012; Matos et al., 2014). We found that there may be shared regulatory logic between CYCD5;1, WEE1, and CYCD6;1/CYCD7;1 in these decisions as well. CYCD6;1 promotes asymmetric divisions of the root cortex-endodermal initials and is a direct target of a complex containing SCR and SHORTROOT (SHR) (Sozzani et al., 2010). The rapid upregulation of the *CYCD6;1* gene in response to SHR induction in roots is well characterized, though less appreciated may be the concomitant upregulation of *WEE1* and *CYCD5;1* (Sozzani et al., 2010). Moreover, *SHR* is required for leaf growth in part through upregulation of genes that promote the G1/S transition, including *CYCD5;1* (Dhondt et al., 2010). In the stomatal lineage, MUTE targets *CYCD5;1*, and *WEE1* is also upregulated upon induced *MUTE* expression (Han et al., 2018). However, the stomatal lineage-restricted *CYCD7;1* is not a target of MUTE (Han et al., 2018; Weimer et al., 2018). *CYCD5* and *WEE1* may thus be induced by core transcriptional regulators to regulate formative divisions, but there are also independent routes of tissue-specific expression, as evidenced by *CYCD7;1*.

It is not clear why certain cell states employ additional core cell cycle regulators. In the stomatal lineage, loss of either *CYCD5;1* or *CYCD7;1* results in the accumulation of larger GMCs and a delay transiting into guard cell fate (Han et al., 2018; Weimer et al., 2018). Loss-of-function *WEE1* phenotypes have only been measured in response to DNA damage in roots, but include a prolonged cell cycle and premature differentiation into vascular cells (Cools et al., 2011). If the role of WEE1 is consistent among cell types, then a plausible outcome for cells expressing both specialized CYCDs and WEE1 would be the production of larger and more transcriptionally active cells, a situation we observed in the profiled stomatal lineage cells, and that may be required during other formative divisions. The cell cycle represents just one regulatory module, and it will be interesting to leverage our developing leaf atlas to examine signaling cascades and complementary modules. Moreover, a comparison of vasculature cell states in the leaf with that of the stem and root may resolve robust programs that control such tissue-specific differentiation both above and below ground, as vasculature threads throughout the plant body. Finally, detecting *WEE1* expression in scRNA-seq data, when it has often been undetectable (Cools et al., 2011; Dhondt et al., 2010), bodes well for being able to identify other conserved modules that evaded previous detection due to low or highly restricted expression patterns.

Single-cell transcriptomics enabled cell states within the developing leaf to be defined by putatively dynamic instances of a given cell transition or identity, corresponding to cell function and lineage relationships (Morris, 2019; Sagar and Grün, 2020; Wagner et al., 2016). This ultimately provides a framework to compare molecular mechanisms of specific cell fate transitions in plant development between species, as well as with animal development. It will be exciting to consider evolutionarily divergent and convergent deployment of specific molecular regulators and developmental strategies. Furthermore, complementary genomics and proteomics approaches will help shape our understanding of underlying cellular modules and programs, such as by providing insight into gene regulatory networks and protein-protein interactions.

## ACKNOWLEDGEMENTS

We are grateful to members of the Bergmann lab for helpful guidance, discussions, and feedback. We also appreciate discussions and feedback from Prof. José Dinneny and his group. Catherine Ballenger provided initial technical assistance; Dr. Andrea Mair and Dr. Ao Liu provided feedback on plant genetics and genomics; Dr. Eva-Sophie Wallner and Dr. Margot Smit provided insight into vasculature development; and Dr. Heather Meyer, Dr. Andrew Muroyama, Dr. Patricia Lang, Gabriel Amador, and Dirk Spencer provided feedback on the manuscript. Cell sorting/flow cytometry analysis was performed at the Stanford Shared FACS Facility. The Stanford Functional Genomics Facility was supported by NIH Instrumentation Grants (S10OD018220 and 1S10OD021763). The Stanford Genomics Center provided computational power and assistance. C.L.A. was supported by an NIH-NIGMS Ruth L. Kirschstein-NRSA postdoctoral fellowship (F32GM129918), A.V. was supported by an EMBO postdoctoral fellowship (ALTF 707-2012), and A.K.W. was supported by a postdoctoral fellowship from Deutsche Forschungsgemeinschaft. D.C.B. is an HHMI investigator.

## AUTHOR CONTRIBUTIONS

C.L.A. conceived, designed, performed all experiments (except as noted below), analyzed results, performed statistical analysis, and wrote the manuscript. A.V. created the *MUTEp::amiSPCH* and *SPCHp::gSPCH-YFP* genomic *SPCH* rescue lines. M.X.A.G. contributed to imaging *SPCH* rescue lines. N.K.S., N.S., and V.G. contributed to creation and imaging of reporter lines. Y.G. performed time-lapse imaging. A.K.W. provided cell cycle insight. D.C.B. analyzed results and wrote the manuscript.

## MATERIALS AND METHODS

### Plant material and growth conditions

All transgenic lines were examined in the *Arabidopsis thaliana* Columbia-0 (Col) ecotype, which served as the wildtype control in all experiments. The following previously described transgenics were used in this study: *ML1p::YFP-RCI2A* (Roeder et al., 2010), *TMMp::TMM-YFP* (Nadeau and Sack, 2002), *SPCHp::cdsSPCH-YFP* and *SPCHp::gSPCH-YFP* (Vatén et al., 2018). Seedlings were grown on half-strength (½) Murashige and Skoog (MS) medium (Caisson Labs MSP01) at 22°C under long-day conditions (16 hr light and 8 hr dark cycles) and were used at the indicated day post germination (dpg) for respective experiments.

### Vector construction and plant transformation

Transgenic promoter reporters fused to a nuclear localized version of YFP (*YFP-NLS*) were generated using the Gateway system (Invitrogen). In brief, promoters were PCR-amplified from genomic DNA and cloned into pENTR then recombined with vectors bearing *YFP-NLS*, into the binary R4pGWB destination vector system (Nakagawa et al., 2008; Tanaka et al., 2011). To generate the *MUTEp::amiSPCH* transgene, the *artificial microRNA* (*amiRNA*) sequence was designed with the Web MicroRNA Designer platform (http://wmd3.weigelworld.org), engineered using the pRS300 plasmid as template, and together with the *MUTE* promoter, was subcloned into the destination vector pGWB. Primer sequences used to generate transgenics are provided in Table S2, and all referenced genes are included in Table S3. Transgenic plants were generated by Agrobacterium-mediated transformation (Clough, 2005; Clough and Bent, 1998), and seedlings were selected on ½ MS plates supplemented with respective antibiotics (e.g. 12 mg/L Basta).

### Analysis of transcriptional and translational reporters

Seedlings from at least two independent lines were collected and analyzed, as indicated for each respective experiment. Fluorescence images were taken with a Leica TCS SP5 microscope (Leica Microsystems) and processed with Fiji (Schindelin et al., 2012). Cell outlines were visualized by either the plasma membrane marker *ML1p:mCherry-RCI2A* or with propidium iodide (Molecular Probes, P3566).

### Quantification of developmental phenotypes

Seedlings from at least two independent and homozygous lines were collected, except for the two independent, segregating lines of *MUTEp::amiSPCH*. Seedlings were cleared in 7:1 ethanol:acetic acid, treated for 30 min with 1 N potassium hydroxide, rinsed in water, and mounted in Hoyer’s medium. Differential contrast interference (DIC) images of the middle region of adaxial epidermis of cotyledons were obtained with a Leica DM2500 microscope (Leica Microsystems).

### Single-cell (sc)RNA-seq and data analysis

#### Single-cell isolation

Single-cells (i.e. protoplasts, plant cells without cell walls) from respective transgenic reporter lines were isolated as described previously (Adrian et al., 2015; Bargmann and Birnbaum, 2010). In brief, whole aerial tissue or first true leaves harvested from ∼180-260 total 10 dpg seedlings were placed into 15 mL protoplasting solution: pH 5.7 with 1 M Tris-Cl pH 7.5, contains 1.25% [w/v] cellulase (Yakult R-10), 0.3% [w/v] macerozyme (Yakult R-10), 0.4 M mannitol (Sigma M1902), 20 mM MES (Caisson Labs M009), 20 mM KCl, 0.1% BSA (Fisher BP1605), and 10 mM CaCl_2_. After 2 hours of gentle shaking, the protoplast solution was filtered through a 40 µm cell strainer (BD Falcon 352340) and centrifuged for 5 min at 500 x g. Protoplast pellets were resuspended in ∼1-3 mL of protoplasting solution and sorted using a FACSAria II (BD Biosciences) instrument fitted with a 100 µm nozzle.

#### Droplet-based 10X Genomics

300,000 single-cell positive events were sorted into ∼1-2mL of protoplasting solution. Kept at room temperature, cells were lightly centrifuged and resuspended in protoplasting solution to yield a sample with ∼2,000 cells/µL in a total volume of ∼20-30 µL. Cell concentration was determined with a hemocytometer, and limited cell lysis was detected by staining with 1:1 trypan blue (Thermo Scientific 12250061). Cells were loaded into a microfluidic device (10X Genomics) with 3’ v2 or v3 chemistry to capture ∼5,000-10,000 cells per sample (Zheng et al., 2017). mRNA was reverse transcribed and cDNA quality was assessed using High Sensitivity NGS Fragment Analyzer (Agilent). Illumina libraries were constructed with Gene Expression v2 and v3 kits (10X Genomics), and sequencing of paired-end 75 bp reads was performed on a HiSeq4000 (Illumina), according to the manufacturer’s instructions. Illumina BCL files were processed via Cell Ranger v2.1.1 and v3.1.0 pipelines, with reads aligned to the Arabidopsis TAIR10 genome assembly (Lamesch et al., 2012). Subsequently, spliced and unspliced transcript matrices were generated using samtools (Li et al., 2009) and velocyto (La Manno et al., 2018).

#### Plate-based Smart-seq2

Single-cell positive events were indexed and sorted directly into single wells of a 96-well plate, which contained 4 µL of lysis buffer with 1 U/µL of Recombinant RNase Inhibitor (RRI, Clontech 2313B), 0.1% [w/v] Triton X-100 (Thermo Scientific 85111), 2.5 mM dNTP (Thermo Scientific 10297018), and 2.5 µM oligodT30VN (5′AAGCAGTGGTATCAACGCAGAGTACT30VN-3′, IDT). Once sorted, cells were immediately spun down and frozen at -80°C for temporary storage. cDNA synthesis was performed as previously described using the Smart-seq2 protocol (Picelli et al., 2014), with SMARTScribe (Clontech 639538) reverse transcriptase and KAPA HiFi DNA polymerase (Kapa Biosystems KK2602). cDNA quality was assessed using High Sensitivity NGS Fragment Analyzer Kit (Advanced Analytical DNF-474), and barcoded Illumina libraries were made using the miniaturized Nextera XT protocol (Mora-Castilla et al., 2016). Sequencing of single-end 75 bp reads was performed on a NextSeq500 (Illumina) high output flow cell, according to the manufacturer’s instructions. FASTQ files were generated from Illumina BCL files and low-quality sequences were trimmed by cutadapt (Marcel, 2011). Reads were aligned to the TAIR10 genome assembly (Lamesch et al., 2012) with STAR (Dobin et al., 2012), and a gene-by-cell raw count matrix was generated by HTSeq (Anders et al., 2015).

#### scRNA-seq dataset analyses

Transcripts Per kilobase Million (TPM) values (Smart-seq2) or gene-barcode matrices (10X Genomics) from single cells were analyzed with Seurat v3 (Butler et al., 2018). Data processing and plotting was done with R/RStudio (R Core Team, 2019; RStudio Team, 2019), except for as noted otherwise. Cells were pre-processed to filter out poor-quality cells with relatively few detected transcripts (i.e. less than 100-500 UMIs/cell), as well as potential cell doublets (i.e. two cells with a shared cell barcode identifier) with relatively high numbers of detected genes. Cells with substantial detected mitochondria and chloroplast transcripts (i.e. more than 25% of UMIs attributed to respective transcripts) were also filtered out, under the assumption that such profiles may represent plastids released from cells.

After pre-processing, the data were log-normalized by NormalizeData and variable genes were identified with FindVariableGenes, with parameters selection.method = “vst” and nfeatures = 2000. For the *TMMp::TMM-YFP* datasets, the n=3 samples from 10X Genomics were integrated before subsequent integration with the Smart-seq2 dataset, using FindIntegrationAnchors and IntegrateData. Unwanted sources of variation derived from cell cycle stage across different tissues were regressed out of the *ML1p::YFP-RCI2A* dataset to avoid clustering based on cell cycle (Tirosh et al., 2016). Cell cycle stage genes were defined by known homology and previous time-course experiments (Menges et al., 2005, 2003; Vandepoele et al., 2002). Representative genes induced during S and G2/M phases (Table S4) were used to assign ‘cell cycle scores’ with CellCycleScoring, which were used in downstream cell cycle regression. The data were then scaled with ScaleData. Principle component analysis (PCA) dimensionality reduction was performed with RunPCA to calculate 50 principal components. A Shared Nearest Neighbor (SNN) graph was generated by FindNeighbors, and graph-based cell clustering based on the SNN was performed by FindClusters. The clustree package (Zappia and Oshlack, 2018) was implemented to visualize how clusters breakdown from one resolution to another. Uniform Manifold Approximation and Projection (UMAP) embedding was performed by RunUMAP, using all 50 principal components with parameters n_neighbors = 30, min_dist = 0.3, umap.method = “uwot”, and metric = “cosine”. Data visualization was performed with ggplot2 (Wickham, 2016).

Differentially expressed genes in each cluster were identified by the non-parametric Wilcoxon rank sum test (default) using Seurat v3 (Butler et al., 2018) FindAllMarkers, with parameters only.pos = “TRUE”, min.pct = 0.25 and logfc.threshold = 0.223 (1.25-fold change). The min.pct argument requires that genes are detected within a minimum percentage (i.e. 25%) of cells of a given cluster, and the logfc.threshold argument requires a minimum fold change (i.e. 1.25-fold) for a gene to be identified as differentially expressed. Enriched GO terms associated with the top 100 differentially expressed genes were identified by clusterProfiler (Yu et al., 2012) compareCluster and gofilter (level = 3) with the following parameters fun = “enrichGO”, OrgDb = “org.At.tair.db”, ont = “BP”, pAdjustMethod = “BH”, pvalueCutoff=0.05, qvalueCutoff=0.05.

Transcriptional dynamics (i.e. predicting future cell states by leveraging unspliced and spliced transcripts) in the *ML1p::YFP-RCI2A* 10X Genomics dataset was assessed by a steady-state deterministic model using scVelo (Bergen et al., 2019), with scv.pp.filter_and_normalize parameters min_shared_counts = 100 and min_counts_u = 500. scVelo was implemented with python using Jupyter/IPython Notebook (Perez and Granger, 2007; Van Rossum and Drake, 2009). Pseudotime lineage inference was performed with the integrated *TMMp::TMM-YFP* datasets in R/RStudio using Slingshot (Street et al., 2018), which constructs a minimum spanning tree and fits simultaneous principle curves through the tree, with the parameter reduceDim = “UMAP”.

#### Accession numbers

The scRNA-seq datasets generated in this study are deposited in GEO under accession number GSXXXXX.

## REFERENCES

Adrian J, Chang J, Ballenger CE, Bargmann BOR, Alassimone J, Davies KA, Lau OS, Matos JL, Hachez C, Lanctot A, Vatén A, Birnbaum KD, Bergmann DC. 2015. Transcriptome dynamics of the stomatal lineage: birth, amplification, and termination of a self-renewing population. Dev Cell 33:107–118. doi: 10.1016/j.devcel.2015.01.025

Albert NW. 2015. Subspecialization of R2R3-MYB Repressors for Anthocyanin and Proanthocyanidin Regulation in Forage Legumes. Front Plant Sci.

Anders S, Pyl PT, Huber W. 2015. HTSeq--a Python framework to work with high-throughput sequencing data. Bioinformatics 31:166–169. doi: 10.1093/bioinformatics/btu638

Bargmann BOR, Birnbaum KD. 2010. Fluorescence activated cell sorting of plant protoplasts. J Vis Exp. doi: 10.3791/1673

Becht E, McInnes L, Healy J, Dutertre C-A, Kwok IWH, Ng LG, Ginhoux F, Newell EW. 2018. Dimensionality reduction for visualizing single-cell data using UMAP. Nat Biotechnol. doi: 10.1038/nbt.4314

Bergen V, Lange M, Peidli S, Wolf FA, Theis FJ. 2019. Generalizing RNA velocity to transient cell states through dynamical modeling. bioRxiv 820936. doi: 10.1101/820936

Bonke M, Thitamadee S, Mähönen AP, Hauser M-T, Helariutta Y. 2003. APL regulates vascular tissue identity in Arabidopsis. Nature 426:181–186. doi: 10.1038/nature02100

Butler A, Hoffman P, Smibert P, Papalexi E, Satija R. 2018. Integrating single-cell transcriptomic data across different conditions, technologies, and species. Nat Biotechnol 36:411–420. doi: 10.1038/nbt.4096

Carlson M. n.d. No Title. doi: 10.18129/B9.bioc.org.At.tair.db

Chen L-Q, Qu X-Q, Hou B-H, Sosso D, Osorio S, Fernie AR, Frommer WB. 2012. Sucrose Efflux Mediated by SWEET Proteins as a Key Step for Phloem Transport. Science (80-) 335:207 LP – 211. doi: 10.1126/science.1213351

Clough SJ. 2005. Floral dip: agrobacterium-mediated germ line transformation. Methods Mol Biol 286:91–102. doi: 10.1385/1-59259-827-7:091

Clough SJ, Bent AF. 1998. Floral dip: a simplified method for Agrobacterium-mediated transformation of Arabidopsis thaliana. Plant J 16:735–743. doi: 10.1046/j.1365-313x.1998.00343.x

Cools T, Iantcheva A, Weimer AK, Boens S, Takahashi N, Maes S, Van den Daele H, Van Isterdael G, Schnittger A, De Veylder L. 2011. The &lt;em&gt;Arabidopsis thaliana&lt;/em&gt; Checkpoint Kinase WEE1 Protects against Premature Vascular Differentiation during Replication Stress. Plant Cell 23:1435 LP – 1448. doi: 10.1105/tpc.110.082768

Cruz-Ramírez A, Díaz-Triviño S, Blilou I, Grieneisen VA, Sozzani R, Zamioudis C, Miskolczi P, Nieuwland J, Benjamins R, Dhonukshe P, Caballero-Pérez J, Horvath B, Long Y, Mähönen AP, Zhang H, Xu J, Murray JAH, Benfey PN, Bako L, Marée AFM, Scheres B. 2012. A bistable circuit involving SCARECROW-RETINOBLASTOMA integrates cues to inform asymmetric stem cell division. Cell 150:1002–1015. doi: 10.1016/j.cell.2012.07.017

Davies KA, Bergmann DC. 2014. Functional specialization of stomatal bHLHs through modification of DNA-binding and phosphoregulation potential. Proc Natl Acad Sci U S A 111:15585–15590. doi: 10.1073/pnas.1411766111

De Rybel B, Möller B, Yoshida S, Grabowicz I, Barbier de Reuille P, Boeren S, Smith RS, Borst JW, Weijers D. 2013. A bHLH complex controls embryonic vascular tissue establishment and indeterminate growth in Arabidopsis. Dev Cell 24:426–437. doi: 10.1016/j.devcel.2012.12.013

De Schutter K, Joubès J, Cools T, Verkest A, Corellou F, Babiychuk E, Van Der Schueren E, Beeckman T, Kushnir S, Inzé D, De Veylder L. 2007. Arabidopsis WEE1 kinase controls cell cycle arrest in response to activation of the DNA integrity checkpoint. Plant Cell 19:211–225. doi: 10.1105/tpc.106.045047

Denyer T, Ma X, Klesen S, Scacchi E, Nieselt K, Timmermans MCP. 2019. Spatiotemporal Developmental Trajectories in the Arabidopsis Root Revealed Using High-Throughput Single-Cell RNA Sequencing. Dev Cell 48:840-852.e5. doi: 10.1016/j.devcel.2019.02.022

Dhondt S, Coppens F, De Winter F, Swarup K, Merks RMH, Inzé D, Bennett MJ, Beemster GTS. 2010. SHORT-ROOT and SCARECROW Regulate Leaf Growth in Arabidopsis by Stimulating S-Phase Progression of the Cell Cycle. Plant Physiol 154:1183 LP – 1195. doi: 10.1104/pp.110.158857

Dobin A, Davis CA, Schlesinger F, Drenkow J, Zaleski C, Jha S, Batut P, Chaisson M, Gingeras TR. 2012. STAR: ultrafast universal RNA-seq aligner. Bioinformatics 29:15–21. doi: 10.1093/bioinformatics/bts635

Durinck S, Moreau Y, Kasprzyk A, Davis S, De Moor B, Brazma A, Huber W. 2005. BioMart and Bioconductor: a powerful link between biological databases and microarray data analysis. Bioinformatics 21:3439–3440. doi: 10.1093/bioinformatics/bti525

Durinck S, Spellman PT, Birney E, Huber W. 2009. Mapping identifiers for the integration of genomic datasets with the R/Bioconductor package biomaRt. Nat Protoc 4:1184–1191. doi: 10.1038/nprot.2009.97

Efroni I, Mello A, Nawy T, Ip P-L, Rahni R, DelRose N, Powers A, Satija R, Birnbaum KD. 2016. Root Regeneration Triggers an Embryo-like Sequence Guided by Hormonal Interactions. Cell 165:1721–1733. doi: 10.1016/j.cell.2016.04.046

Fisher K, Turner S. 2007. PXY, a receptor-like kinase essential for maintaining polarity during plant vascular-tissue development. Curr Biol 17:1061–1066. doi: 10.1016/j.cub.2007.05.049

Gulati GS, Sikandar SS, Wesche DJ, Manjunath A, Bharadwaj A, Berger MJ, Ilagan F, Kuo AH, Hsieh RW, Cai S, Zabala M, Scheeren FA, Lobo NA, Qian D, Yu FB, Dirbas FM, Clarke MF, Newman AM. 2020. Single-cell transcriptional diversity is a hallmark of developmental potential. Science (80-) 367:405 LP – 411. doi: 10.1126/science.aax0249

Gutierrez C. 2016. 25 Years of Cell Cycle Research: What’s Ahead? Trends Plant Sci 21:823–833. doi: 10.1016/j.tplants.2016.06.007

Han C, Liu Y, Shi W, Qiao Y, Wang L, Tian Y, Fan M, Deng Z, Lau OS, De Jaeger G, Bai M-Y. 2020. KIN10 promotes stomatal development through stabilization of the SPEECHLESS transcription factor. Nat Commun 11:4214. doi: 10.1038/s41467-020-18048-w

Han S-K, Qi X, Sugihara K, Dang JH, Endo TA, Miller KL, Kim E-D, Miura T, Torii KU. 2018. MUTE Directly Orchestrates Cell-State Switch and the Single Symmetric Division to Create Stomata. Dev Cell 45:303-315.e5. doi: 10.1016/j.devcel.2018.04.010

Hara K, Kajita R, Torii KU, Bergmann DC, Kakimoto T. 2007. The secretory peptide gene EPF1 enforces the stomatal one-cell-spacing rule. Genes Dev 21:1720–1725. doi: 10.1101/gad.1550707

Hara K, Yokoo T, Kajita R, Onishi T, Yahata S, Peterson KM, Torii KU, Kakimoto T. 2009. Epidermal cell density is autoregulated via a secretory peptide, EPIDERMAL PATTERNING FACTOR 2 in Arabidopsis leaves. Plant Cell Physiol 50:1019–1031. doi: 10.1093/pcp/pcp068

Hellmann E, Ko D, Ruonala R, Helariutta Y. 2018. Plant Vascular Tissues-Connecting Tissue Comes in All Shapes. Plants (Basel, Switzerland) 7:109. doi: 10.3390/plants7040109

Hetherington AM, Woodward FI. 2003. The role of stomata in sensing and driving environmental change. Nature 424:901–908. doi: 10.1038/nature01843

Hunt L, Bailey KJ, Gray JE. 2010. The signalling peptide EPFL9 is a positive regulator of stomatal development. New Phytol 186:609–614. doi: 10.1111/j.1469-8137.2010.03200.x

Iida H, Yoshida A, Takada S. 2019. ATML1 activity is restricted to the outermost cells of the embryo through post-transcriptional repressions. Dev 146:dev169300. doi: 10.1242/dev.169300

Kanaoka MM, Pillitteri LJ, Fujii H, Yoshida Y, Bogenschutz NL, Takabayashi J, Zhu J-K, Torii KU. 2008. SCREAM/ICE1 and SCREAM2 specify three cell-state transitional steps leading to arabidopsis stomatal differentiation. Plant Cell 20:1775–1785. doi: 10.1105/tpc.108.060848

Karaiskos N, Wahle P, Alles J, Boltengagen A, Ayoub S, Kipar C, Kocks C, Rajewsky N, Zinzen RP. 2017. The Drosophila embryo at single-cell transcriptome resolution. Science (80-) 358:194 LP – 199. doi: 10.1126/science.aan3235

Kidner CA, Timmermans MCP. 2010. Signaling sides adaxial-abaxial patterning in leaves. Curr Top Dev Biol 91:141–168. doi: 10.1016/S0070-2153(10)91005-3

Kondo T, Kajita R, Miyazaki A, Hokoyama M, Nakamura-Miura T, Mizuno S, Masuda Y, Irie K, Tanaka Y, Takada S, Kakimoto T, Sakagami Y. 2010. Stomatal density is controlled by a mesophyll-derived signaling molecule. Plant Cell Physiol 51:1–8. doi: 10.1093/pcp/pcp180

Kubo M, Udagawa M, Nishikubo N, Horiguchi G, Yamaguchi M, Ito J, Mimura T, Fukuda H, Demura T. 2005. Transcription switches for protoxylem and metaxylem vessel formation. Genes Dev 19:1855–1860. doi: 10.1101/gad.1331305

Kumari A, Jewaria PK, Bergmann DC, Kakimoto T. 2014. Arabidopsis reduces growth under osmotic stress by decreasing SPEECHLESS protein. Plant Cell Physiol 55:2037–2046. doi: 10.1093/pcp/pcu159

La Manno G, Soldatov R, Zeisel A, Braun E, Hochgerner H, Petukhov V, Lidschreiber K, Kastriti ME, Lönnerberg P, Furlan A, Fan J, Borm LE, Liu Z, van Bruggen D, Guo J, He X, Barker R, Sundström E, Castelo-Branco G, Cramer P, Adameyko I, Linnarsson S, Kharchenko P V. 2018. RNA velocity of single cells. Nature 560:494–498. doi: 10.1038/s41586-018-0414-6

Lai LB, Nadeau JA, Lucas J, Lee E-K, Nakagawa T, Zhao L, Geisler M, Sack FD. 2005. The Arabidopsis R2R3 MYB proteins FOUR LIPS and MYB88 restrict divisions late in the stomatal cell lineage. Plant Cell 17:2754–2767. doi: 10.1105/tpc.105.034116

Lamesch P, Berardini TZ, Li D, Swarbreck D, Wilks C, Sasidharan R, Muller R, Dreher K, Alexander DL, Garcia-Hernandez M, Karthikeyan AS, Lee CH, Nelson WD, Ploetz L, Singh S, Wensel A, Huala E. 2012. The Arabidopsis Information Resource (TAIR): improved gene annotation and new tools. Nucleic Acids Res 40:D1202–D1210. doi: 10.1093/nar/gkr1090

Lampard GR, MacAlister CA, Bergmann DC. 2008. &lt;em&gt;Arabidopsis&lt;/em&gt; Stomatal Initiation Is Controlled by MAPK-Mediated Regulation of the bHLH SPEECHLESS. Science (80-) 322:1113 LP – 1116. doi: 10.1126/science.1162263

Lau OS, Davies KA, Chang J, Adrian J, Rowe MH, Ballenger CE, Bergmann DC. 2014. Direct roles of SPEECHLESS in the specification of stomatal self-renewing cells. Science 345:1605–1609. doi: 10.1126/science.1256888

Lau OS, Song Z, Zhou Z, Davies KA, Chang J, Yang X, Wang S, Lucyshyn D, Tay IHZ, Wigge PA, Bergmann DC. 2018. Direct Control of SPEECHLESS by PIF4 in the High-Temperature Response of Stomatal Development. Curr Biol 28:1273-1280.e3. doi: 10.1016/j.cub.2018.02.054

Lauber MH, Waizenegger I, Steinmann T, Schwarz H, Mayer U, Hwang I, Lukowitz W, Jürgens G. 1997. The Arabidopsis KNOLLE protein is a cytokinesis-specific syntaxin. J Cell Biol 139:1485–1493. doi: 10.1083/jcb.139.6.1485

Lee LR, Bergmann DC. 2019. The plant stomatal lineage at a glance. J Cell Sci 132. doi: 10.1242/jcs.228551

Lee LR, Wengier DL, Bergmann DC. 2019. Cell-type-specific transcriptome and histone modification dynamics during cellular reprogramming in the Arabidopsis stomatal lineage. Proc Natl Acad Sci U S A 116:21914–21924. doi: 10.1073/pnas.1911400116

Li H, Handsaker B, Wysoker A, Fennell T, Ruan J, Homer N, Marth G, Abecasis G, Durbin R. 2009. The Sequence Alignment/Map format and SAMtools. Bioinformatics 25:2078–2079. doi: 10.1093/bioinformatics/btp352

Lu P, Porat R, Nadeau JA, O’Neill SD. 1996. Identification of a meristem L1 layer-specific gene in Arabidopsis that is expressed during embryonic pattern formation and defines a new class of homeobox genes. Plant Cell 8:2155 LP – 2168. doi: 10.1105/tpc.8.12.2155

Lukowitz W, Mayer U, Jürgens G. 1996. Cytokinesis in the Arabidopsis embryo involves the syntaxin-related KNOLLE gene product. Cell 84:61–71. doi: 10.1016/s0092-8674(00)80993-9

MacAlister CA, Ohashi-Ito K, Bergmann DC. 2007. Transcription factor control of asymmetric cell divisions that establish the stomatal lineage. Nature 445:537–540. doi: 10.1038/nature05491

Marcel M. 2011. Cutadapt removes adapter sequences from high-throughput sequencing reads. EMBnet J 17:10–12.

Matos JL, Lau OS, Hachez C, Cruz-Ramírez A, Scheres B, Bergmann DC. 2014. Irreversible fate commitment in the Arabidopsis stomatal lineage requires a FAMA and RETINOBLASTOMA-RELATED module. Elife 3:1–15. doi: 10.7554/eLife.03271

McInnes L, Healy J, Saul N, Grossberger L. 2018. UMAP: Uniform Manifold Approximation and Projection. J Open Source Softw 3:861.

Menges M, de Jager SM, Gruissem W, Murray JAH. 2005. Global analysis of the core cell cycle regulators of Arabidopsis identifies novel genes, reveals multiple and highly specific profiles of expression and provides a coherent model for plant cell cycle control. Plant J 41:546–566. doi: 10.1111/j.1365-313X.2004.02319.x

Menges M, Hennig L, Gruissem W, Murray JAH. 2003. Genome-wide gene expression in an Arabidopsis cell suspension. Plant Mol Biol 53:423–442. doi: 10.1023/B:PLAN.0000019059.56489.ca

Mora-Castilla S, To C, Vaezeslami S, Morey R, Srinivasan S, Dumdie JN, Cook-Andersen H, Jenkins J, Laurent LC. 2016. Miniaturization Technologies for Efficient Single-Cell Library Preparation for Next-Generation Sequencing. J Lab Autom 21:557–567. doi: 10.1177/2211068216630741

Morris SA. 2019. The evolving concept of cell identity in the single cell era. Development 146:dev169748. doi: 10.1242/dev.169748

Mott KA. 2007. Leaf hydraulic conductivity and stomatal responses to humidity in amphistomatous leaves. Plant Cell Environ 30:1444–1449. doi: 10.1111/j.1365-3040.2007.01720.x

Nadeau JA, Sack FD. 2002. Control of stomatal distribution on the Arabidopsis leaf surface. Science 296:1697–1700. doi: 10.1126/science.1069596

Nakagawa T, Nakamura S, Tanaka K, Kawamukai M, Suzuki T, Nakamura K, Kimura T, Ishiguro S. 2008. Development of R4 gateway binary vectors (R4pGWB) enabling high-throughput promoter swapping for plant research. Biosci Biotechnol Biochem 72:624–629. doi: 10.1271/bbb.70678

Negi J, Moriwaki K, Konishi M, Yokoyama R, Nakano T, Kusumi K, Hashimoto-Sugimoto M, Schroeder JI, Nishitani K, Yanagisawa S, Iba K. 2013. A Dof transcription factor, SCAP1, is essential for the development of functional stomata in Arabidopsis. Curr Biol 23:479–484. doi: 10.1016/j.cub.2013.02.001

Nelms B, Walbot V. 2019. Defining the developmental program leading to meiosis in maize. Science 364:52–56. doi: 10.1126/science.aav6428

Nurse P, Thuriaux P. 1980. Regulatory genes controlling mitosis in the fission yeast Schizosaccharomyces pombe. Genetics 96:627–637.

Nylander M, Heino P, Helenius E, Tapio Palva E, Ronne H, Welin B V. 2001. The low-temperature- and salt-induced RCI2A gene of Arabidopsis complements the sodium sensitivity caused by a deletion of the homologous yeast gene SNA1. Plant Mol Biol 45:341–352. doi: 10.1023/A:1006451914231

Ohashi-Ito K, Bergmann DC. 2006. Arabidopsis FAMA controls the final proliferation/differentiation switch during stomatal development. Plant Cell 18:2493–2505. doi: 10.1105/tpc.106.046136

Perez F, Granger BE. 2007. IPython: A System for Interactive Scientific Computing. Comput Sci Engg 9:21–29. doi: 10.1109/MCSE.2007.53

Picelli S, Faridani OR, Björklund AK, Winberg G, Sagasser S, Sandberg R. 2014. Full-length RNA-seq from single cells using Smart-seq2. Nat Protoc 9:171–181. doi: 10.1038/nprot.2014.006

Pillitteri LJ, Sloan DB, Bogenschutz NL, Torii KU. 2007. Termination of asymmetric cell division and differentiation of stomata. Nature 445:501–505. doi: 10.1038/nature05467

Pillitteri LJ, Torii KU. 2012. Mechanisms of stomatal development. Annu Rev Plant Biol 63:591–614. doi: 10.1146/annurev-arplant-042811-105451

Pruitt RE, Vielle-Calzada JP, Ploense SE, Grossniklaus U, Lolle SJ. 2000. FIDDLEHEAD, a gene required to suppress epidermal cell interactions in Arabidopsis, encodes a putative lipid biosynthetic enzyme. Proc Natl Acad Sci U S A 97:1311–1316. doi: 10.1073/pnas.97.3.1311

Putarjunan A, Ruble J, Srivastava A, Zhao C, Rychel AL, Hofstetter AK, Tang X, Zhu J-K, Tama F, Zheng N, Torii KU. 2019. Bipartite anchoring of SCREAM enforces stomatal initiation by coupling MAP kinases to SPEECHLESS. Nat plants 5:742–754. doi: 10.1038/s41477-019-0440-x

R Core Team. 2019. R: A Language and Environment for Statistical Computing.

Ram H, Sahadevan S, Gale N, Caggiano MP, Yu X, Ohno C, Heisler MG. 2020. An integrated analysis of cell-type specific gene expression reveals genes regulated by REVOLUTA and KANADI1 in the Arabidopsis shoot apical meristem. PLOS Genet 16:e1008661.

Roeder AHK, Chickarmane V, Cunha A, Obara B, Manjunath BS, Meyerowitz EM. 2010. Variability in the control of cell division underlies sepal epidermal patterning in Arabidopsis thaliana. PLoS Biol 8:e1000367. doi: 10.1371/journal.pbio.1000367

RStudio Team. 2019. RStudio: Integrated Development for R.

Ryu KH, Huang L, Kang HM, Schiefelbein J. 2019. Single-Cell RNA Sequencing Resolves Molecular Relationships Among Individual Plant Cells. Plant Physiol 179:1444–1456. doi: 10.1104/pp.18.01482

Saelens W, Cannoodt R, Todorov H, Saeys Y. 2019. A comparison of single-cell trajectory inference methods. Nat Biotechnol 37:547–554. doi: 10.1038/s41587-019-0071-9

Sagar, Grün D. 2020. Deciphering Cell Fate Decision by Integrated Single-Cell Sequencing Analysis. Annu Rev Biomed Data Sci 3:1–22. doi: 10.1146/annurev-biodatasci-111419-091750

Sawchuk MG, Donner TJ, Head P, Scarpella E. 2008. Unique and Overlapping Expression Patterns among Members of Photosynthesis-Associated Nuclear Gene Families in Arabidopsis. Plant Physiol 148:1908–1924. doi: 10.1104/pp.108.126946

Scarpella E, Francis P, Berleth T. 2004. Stage-specific markers define early steps of procambium development in Arabidopsis leaves and correlate termination of vein formation with mesophyll differentiation. Development 131:3445–3455. doi: 10.1242/dev.01182

Schaum N, Karkanias J, Neff NF, May AP, Quake SR, Wyss-Coray T, Darmanis S, Batson J, Botvinnik O, Chen MB, Chen S, Green F, Jones RC, Maynard A, Penland L, Pisco AO, Sit R V, Stanley GM, Webber JT, Zanini F, Baghel AS, Bakerman I, Bansal I, Berdnik D, Bilen B, Brownfield D, Cain C, Chen MB, Chen S, Cho M, Cirolia G, Conley SD, Darmanis S, Demers A, Demir K, de Morree A, Divita T, du Bois H, Dulgeroff LBT, Ebadi H, Espinoza FH, Fish M, Gan Q, George BM, Gillich A, Green F, Genetiano G, Gu X, Gulati GS, Hang Y, Hosseinzadeh S, Huang A, Iram T, Isobe T, Ives F, Jones RC, Kao KS, Karnam G, Kershner AM, Kiss BM, Kong W, Kumar ME, Lam JY, Lee DP, Lee SE, Li G, Li Q, Liu L, Lo A, Lu W-J, Manjunath A, May AP, May KL, May OL, Maynard A, McKay M, Metzger RJ, Mignardi M, Min D, Nabhan AN, Neff NF, Ng KM, Noh J, Patkar R, Peng WC, Penland L, Puccinelli R, Rulifson EJ, Schaum N, Sikandar SS, Sinha R, Sit R V, Szade K, Tan W, Tato C, Tellez K, Travaglini KJ, Tropini C, Waldburger L, van Weele LJ, Wosczyna MN, Xiang J, Xue S, Youngyunpipatkul J, Zanini F, Zardeneta ME, Zhang F, Zhou L, Bansal I, Chen S, Cho M, Cirolia G, Darmanis S, Demers A, Divita T, Ebadi H, Genetiano G, Green F, Hosseinzadeh S, Ives F, Lo A, May AP, Maynard A, McKay M, Neff NF, Penland L, Sit R V, Tan W, Waldburger L, Youngyunpipatkul J, Batson J, Botvinnik O, Castro P, Croote D, Darmanis S, DeRisi JL, Karkanias J, Pisco AO, Stanley GM, Webber JT, Zanini F, Baghel AS, Bakerman I, Batson J, Bilen B, Botvinnik O, Brownfield D, Chen MB, Darmanis S, Demir K, de Morree A, Ebadi H, Espinoza FH, Fish M, Gan Q, George BM, Gillich A, Gu X, Gulati GS, Hang Y, Huang A, Iram T, Isobe T, Karnam G, Kershner AM, Kiss BM, Kong W, Kuo CS, Lam JY, Lehallier B, Li G, Li Q, Liu L, Lu W-J, Min D, Nabhan AN, Ng KM, Nguyen PK, Patkar R, Peng WC, Penland L, Rulifson EJ, Schaum N, Sikandar SS, Sinha R, Szade K, Tan SY, Tellez K, Travaglini KJ, Tropini C, van Weele LJ, Wang BM, Wosczyna MN, Xiang J, Yousef H, Zhou L, Batson J, Botvinnik O, Chen S, Darmanis S, Green F, May AP, Maynard A, Pisco AO, Quake SR, Schaum N, Stanley GM, Webber JT, Wyss-Coray T, Zanini F, Beachy PA, Chan CKF, de Morree A, George BM, Gulati GS, Hang Y, Huang KC, Iram T, Isobe T, Kershner AM, Kiss BM, Kong W, Li G, Li Q, Liu L, Lu W-J, Nabhan AN, Ng KM, Nguyen PK, Peng WC, Rulifson EJ, Schaum N, Sikandar SS, Sinha R, Szade K, Travaglini KJ, Tropini C, Wang BM, Weinberg K, Wosczyna MN, Wu SM, Yousef H, Barres BA, Beachy PA, Chan CKF, Clarke MF, Darmanis S, Huang KC, Karkanias J, Kim SK, Krasnow MA, Kumar ME, Kuo CS, May AP, Metzger RJ, Neff NF, Nusse R, Nguyen PK, Rando TA, Sonnenburg J, Wang BM, Weinberg K, Weissman IL, Wu SM, Quake SR, Wyss-Coray T, Consortium TTM, coordination O, coordination L, processing O collection and, sequencing L preparation and, analysis C data, annotation C type, group W, group S text writing, investigators P. 2018. Single-cell transcriptomics of 20 mouse organs creates a Tabula Muris. Nature 562:367–372. doi: 10.1038/s41586-018-0590-4

Schindelin J, Arganda-Carreras I, Frise E, Kaynig V, Longair M, Pietzsch T, Preibisch S, Rueden C, Saalfeld S, Schmid B, Tinevez J-Y, White DJ, Hartenstein V, Eliceiri K, Tomancak P, Cardona A. 2012. Fiji: an open-source platform for biological-image analysis. Nat Methods 9:676–682. doi: 10.1038/nmeth.2019

Sebé-Pedrós A, Saudemont B, Chomsky E, Plessier F, Mailhé M-P, Renno J, Loe-Mie Y, Lifshitz A, Mukamel Z, Schmutz S, Novault S, Steinmetz PRH, Spitz F, Tanay A, Marlow H. 2018. Cnidarian Cell Type Diversity and Regulation Revealed by Whole-Organism Single-Cell RNA-Seq. Cell 173:1520-1534.e20. doi: 10.1016/j.cell.2018.05.019

Shahan R, Hsu C-W, Nolan TM, + BJC, Taylor IW, Hendrika A, Vlot C, Benfey PN, Ohler U. 2020. A single cell Arabidopsis root atlas reveals developmental trajectories in wild type and cell identity mutants. bioRxiv 2020.06.29.178863. doi: 10.1101/2020.06.29.178863

Shulse CN, Cole BJ, Ciobanu D, Lin J, Yoshinaga Y, Gouran M, Turco GM, Zhu Y, O’Malley RC, Brady SM, Dickel DE. 2019. High-Throughput Single-Cell Transcriptome Profiling of Plant Cell Types. Cell Rep 27:2241-2247.e4. doi: 10.1016/j.celrep.2019.04.054

Siebert S, Farrell JA, Cazet JF, Abeykoon Y, Primack AS, Schnitzler CE, Juliano CE. 2019. Stem cell differentiation trajectories in Hydra resolved at single-cell resolution. Science (80-) 365:eaav9314. doi: 10.1126/science.aav9314

Sorrell DA, Marchbank A, McMahon K, Dickinson JR, Rogers HJ, Francis D. 2002. A WEE1 homologue from Arabidopsis thaliana. Planta 215:518–522. doi: 10.1007/s00425-002-0815-4

Sozzani R, Cui H, Moreno-Risueno MA, Busch W, Van Norman JM, Vernoux T, Brady SM, Dewitte W, Murray JAH, Benfey PN. 2010. Spatiotemporal regulation of cell-cycle genes by SHORTROOT links patterning and growth. Nature 466:128–132. doi: 10.1038/nature09143

Street K, Risso D, Fletcher RB, Das D, Ngai J, Yosef N, Purdom E, Dudoit S. 2018. Slingshot: cell lineage and pseudotime inference for single-cell transcriptomics. BMC Genomics 19:477. doi: 10.1186/s12864-018-4772-0

Sugano SS, Shimada T, Imai Y, Okawa K, Tamai A, Mori M, Hara-Nishimura I. 2010. Stomagen positively regulates stomatal density in Arabidopsis. Nature 463:241–244. doi: 10.1038/nature08682

Takahashi I, Kojima S, Sakaguchi N, Umeda-Hara C, Umeda M. 2010. Two Arabidopsis cyclin A3s possess G1 cyclin-like features. Plant Cell Rep 29:307–315. doi: 10.1007/s00299-010-0817-9

Tanaka Y, Nakamura S, Kawamukai M, Koizumi N, Nakagawa T. 2011. Development of a series of gateway binary vectors possessing a tunicamycin resistance gene as a marker for the transformation of Arabidopsis thaliana. Biosci Biotechnol Biochem 75:804–807. doi: 10.1271/bbb.110063

Tian C, Wang Y, Yu H, He J, Wang J, Shi B, Du Q, Provart NJ, Meyerowitz EM, Jiao Y. 2019. A gene expression map of shoot domains reveals regulatory mechanisms. Nat Commun 10:141. doi: 10.1038/s41467-018-08083-z

Tintori SC, Osborne Nishimura E, Golden P, Lieb JD, Goldstein B. 2016. A Transcriptional Lineage of the Early C. elegans Embryo. Dev Cell 38:430–444. doi: 10.1016/j.devcel.2016.07.025

Tirosh I, Izar B, Prakadan SM, Wadsworth MH, Treacy D, Trombetta JJ, Rotem A, Rodman C, Lian C, Murphy G, Fallahi-Sichani M, Dutton-Regester K, Lin J-R, Cohen O, Shah P, Lu D, Genshaft AS, Hughes TK, Ziegler CGK, Kazer SW, Gaillard A, Kolb KE, Villani A-C, Johannessen CM, Andreev AY, Van Allen EM, Bertagnolli M, Sorger PK, Sullivan RJ, Flaherty KT, Frederick DT, Jané-Valbuena J, Yoon CH, Rozenblatt-Rosen O, Shalek AK, Regev A, Garraway LA. 2016. Dissecting the multicellular ecosystem of metastatic melanoma by single-cell RNA-seq. Science (80-) 352:189 LP – 196. doi: 10.1126/science.aad0501

Triviño M, Martín-Trillo M, Ballesteros I, Delgado D, de Marcos A, Desvoyes B, Gutiérrez C, Mena M, Fenoll C. 2013. Timely expression of the Arabidopsis stoma-fate master regulator MUTE is required for specification of other epidermal cell types. Plant J 75:808–822. doi: 10.1111/tpj.12244

Van Rossum G, Drake FL. 2009. Python 3 Reference Manual. Scotts Valley, CA: CreateSpace.

Vandepoele K, Raes J, De Veylder L, Rouzé P, Rombauts S, Inzé D. 2002. Genome-wide analysis of core cell cycle genes in Arabidopsis. Plant Cell 14:903–916. doi: 10.1105/tpc.010445

Vatén A, Soyars CL, Tarr PT, Nimchuk ZL, Bergmann DC. 2018. Modulation of Asymmetric Division Diversity through Cytokinin and SPEECHLESS Regulatory Interactions in the Arabidopsis Stomatal Lineage. Dev Cell 47:53-66.e5. doi: 10.1016/j.devcel.2018.08.007

von Groll U, Berger D, Altmann T. 2002. The Subtilisin-Like Serine Protease SDD1 Mediates Cell-to-Cell Signaling during Arabidopsis Stomatal Development. Plant Cell 14:1527 LP – 1539. doi: 10.1105/tpc.001016

Wagner A, Regev A, Yosef N. 2016. Revealing the vectors of cellular identity with single-cell genomics. Nat Biotechnol 34:1145–1160. doi: 10.1038/nbt.3711

Wagner DE, Weinreb C, Collins ZM, Briggs JA, Megason SG, Klein AM. 2018. Single-cell mapping of gene expression landscapes and lineage in the zebrafish embryo. Science (80-) 360:981 LP – 987. doi: 10.1126/science.aar4362

Wallner E-S, López-Salmerón V, Belevich I, Poschet G, Jung I, Grünwald K, Sevilem I, Jokitalo E, Hell R, Helariutta Y, Agustí J, Lebovka I, Greb T. 2017. Strigolactone- and Karrikin-Independent SMXL Proteins Are Central Regulators of Phloem Formation. Curr Biol 27:1241–1247. doi: 10.1016/j.cub.2017.03.014

Weimer AK, Matos JL, Sharma N, Patell F, Murray JAH, Dewitte W, Bergmann DC. 2018. Lineage- and stage-specific expressed CYCD7;1 coordinates the single symmetric division that creates stomatal guard cells. Development 145:dev160671. doi: 10.1242/dev.160671

Wickham H. 2016. ggplot2: Elegant Graphics for Data Analysis. Springer-Verlag New York.

Yadav RK, Tavakkoli M, Xie M, Girke T, Reddy GV. 2014. A high-resolution gene expression map of the Arabidopsis shoot meristem stem cell niche. Development 141:2735–2744. doi: 10.1242/dev.106104

Yephremov A, Wisman E, Huijser P, Huijser C, Wellesen K, Saedler H. 1999. Characterization of the FIDDLEHEAD gene of Arabidopsis reveals a link between adhesion response and cell differentiation in the epidermis. Plant Cell 11:2187–2201. doi: 10.1105/tpc.11.11.2187

Yu G, Wang L-G, Han Y, He Q-Y. 2012. clusterProfiler: an R package for comparing biological themes among gene clusters. OMICS 16:284–287. doi: 10.1089/omi.2011.0118

Zappia L, Oshlack A. 2018. Clustering trees: a visualization for evaluating clusterings at multiple resolutions. Gigascience 7. doi: 10.1093/gigascience/giy083

Zhang T-Q, Xu Z-G, Shang G-D, Wang J-W. 2019. A Single-Cell RNA Sequencing Profiles the Developmental Landscape of Arabidopsis Root. Mol Plant 12:648–660. doi: 10.1016/j.molp.2019.04.004

Zheng GXY, Terry JM, Belgrader P, Ryvkin P, Bent ZW, Wilson R, Ziraldo SB, Wheeler TD, McDermott GP, Zhu J, Gregory MT, Shuga J, Montesclaros L, Underwood JG, Masquelier DA, Nishimura SY, Schnall-Levin M, Wyatt PW, Hindson CM, Bharadwaj R, Wong A, Ness KD, Beppu LW, Deeg HJ, McFarland C, Loeb KR, Valente WJ, Ericson NG, Stevens EA, Radich JP, Mikkelsen TS, Hindson BJ, Bielas JH. 2017. Massively parallel digital transcriptional profiling of single cells. Nat Commun 8:14049. doi: 10.1038/ncomms14049

