## Supplemental Files S1-9 for "Single-Cell Resolution of Lineage Trajectories in the Arabidopsis Stomatal Lineage and Developing Leaf"

### SUPPLEMENTARY FIGURES

**Figure S1**. **Leaf atlas experimental workflow and dataset attributes**

**Figure S2**. **Clustering analysis reveals similarities between clusters**

**Figure S3**. **The leaf atlas predicted onset of gene promoter activity within epidermal tissue**

**Figure S4**. **Polarity signatures within the leaf atlas**

**Figure S5**. **Distinct and overlapping cell cycle signatures**

**Figure S6**. **Experimental workflow and dataset attributes for stomatal lineage profiles**

**Figure S7**. **Clustering analysis reveals similarities between clusters**

**Figure S8**. **Onset of predicted gene promoter activity and expression within the stomatal lineage**

**Figure S9**. **The stomatal lineage requires fine-tuned *SPCH* gene expression**

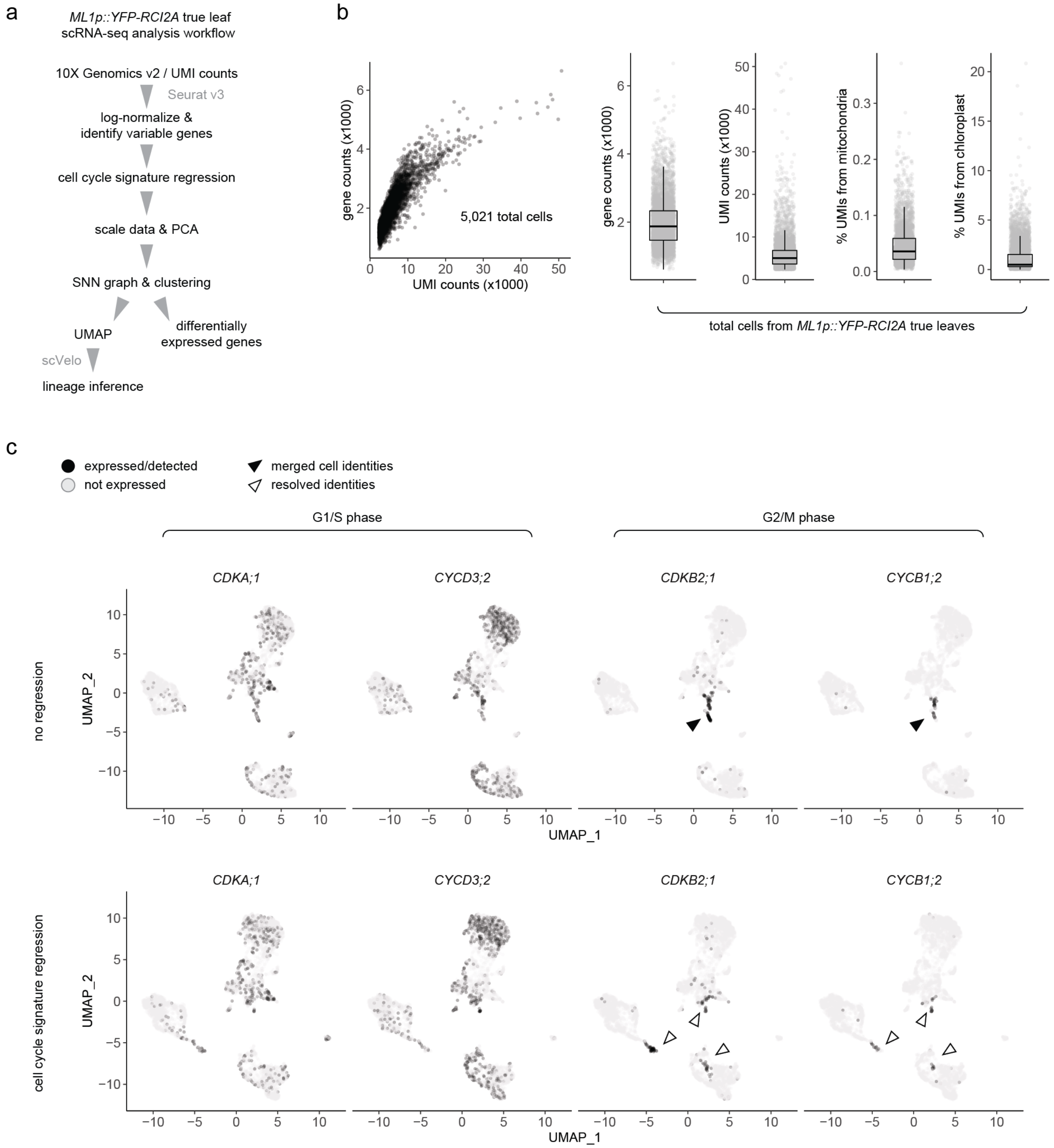


**Figure S1**. **Leaf atlas experimental workflow and dataset attributes.** (a) Workflow implemented to analyze the 10X Genomics dataset from developing shoot tissue of 10 dpg *ML1p::YFP-RCI2A* seedlings. (b) Dataset attributes indicate the number of gene and UMI counts per cell (scatter plot, left; boxplots, right), as well as the percent of UMIs from the mitochondria or chloroplast (boxplots, right), which altogether serve as indicators of sample quality. (c) Representative cell cycle gene expression profiles (black: expressed/detected, grey: not expressed/detected) compared between distinct UMAP plots performed without (top row) and with (bottom row) cell cycle regression. Without regression, cell cycle gene signature masked/merged cell identities (black arrowhead), but dividing cells within different tissues could be resolved after regression (outlined arrowheads).

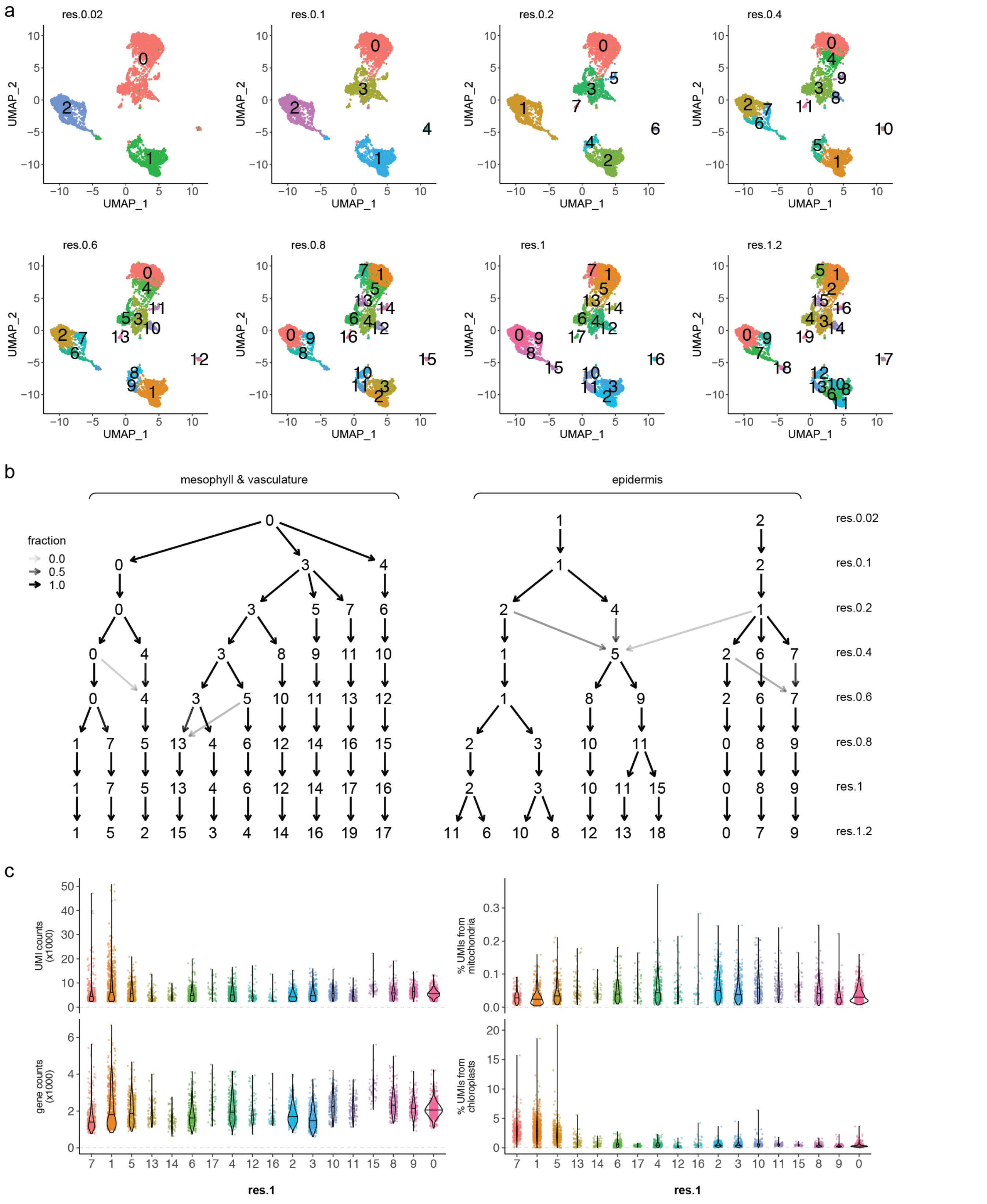

**Figure S2**. **Clustering analysis reveals similarities between clusters.** (a) A range of clustering resolutions depicted on UMAP plots of the developing leaf atlas. (b) Clustering analysis with clustree depicts the cluster breakdown from one resolution to another, with larger clusters divided into smaller clusters at more refined resolutions. Each resolution is on a given ‘row’ in the diagram, with broader to more refined resolutions plotted from top to bottom. Annotated are clusters that correspond to mesophyll and vasculature inner tissue, as well as the epidermis. (c) Dataset attributes within clusters from resolution 1. Though some variation is expected from cluster to cluster, those annotated as mesophyll (clusters 7, 1, and 5) accordingly exhibit a higher percentage of UMIs from chloroplasts.

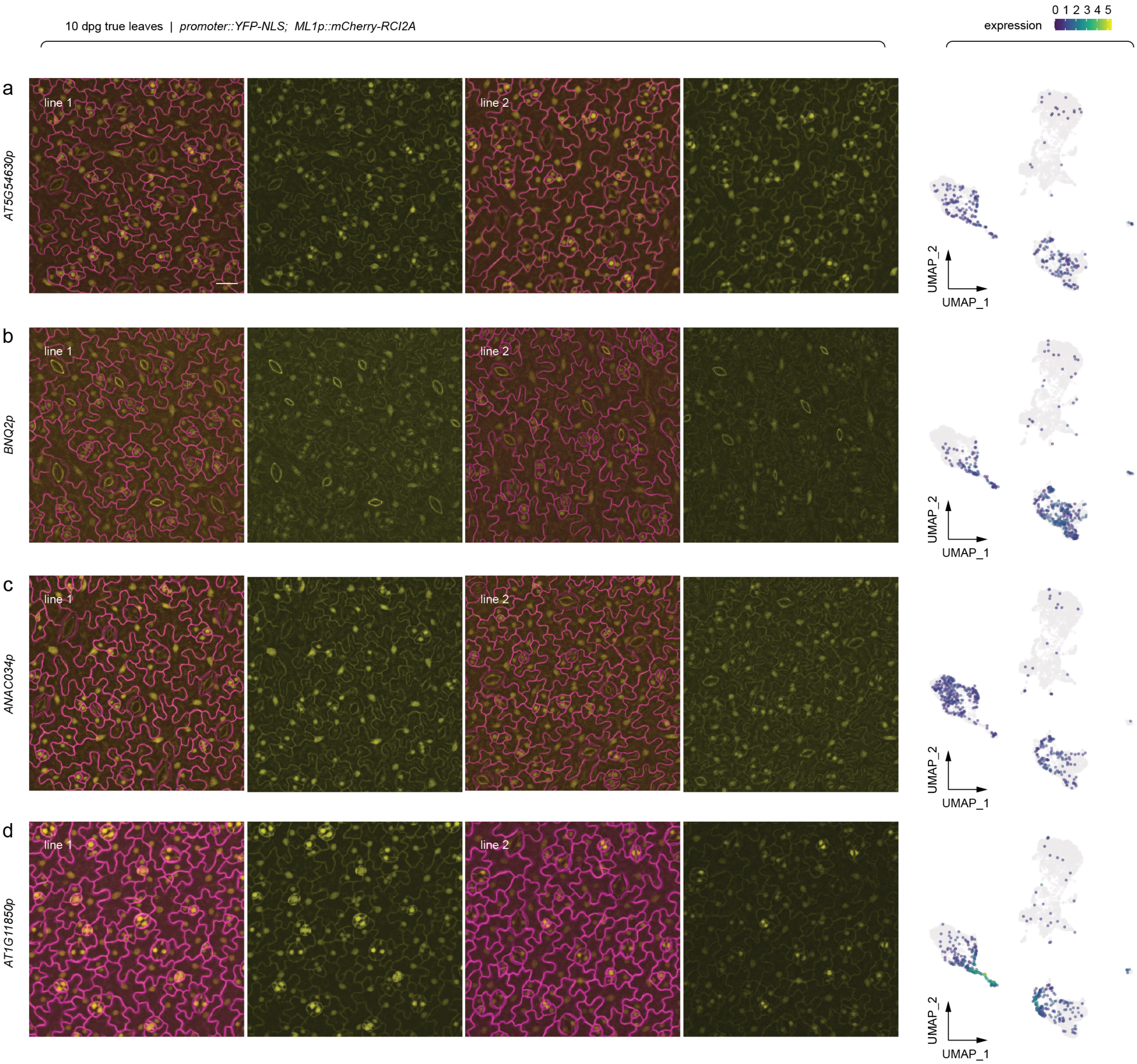


**Figure S3. The leaf atlas predicted onset of gene promoter activity within epidermal tissue.** Representative confocal images of abaxial epidermis (first true leaves) from 10 dpg seedlings of two independent reporter lines. Expression of *YFP-NLS* reporters were driven by respective promoters, and cell outlines were visualized by *ML1p::mCherry-RCI2A*. The same field with and without the *ML1p::mCherry-RCI2A* channel is presented for each reporter. Predicted scRNA-seq expression profiles are depicted in the corresponding UMAP plots. All images were taken at the same magnification (scale bar: 20 µm).

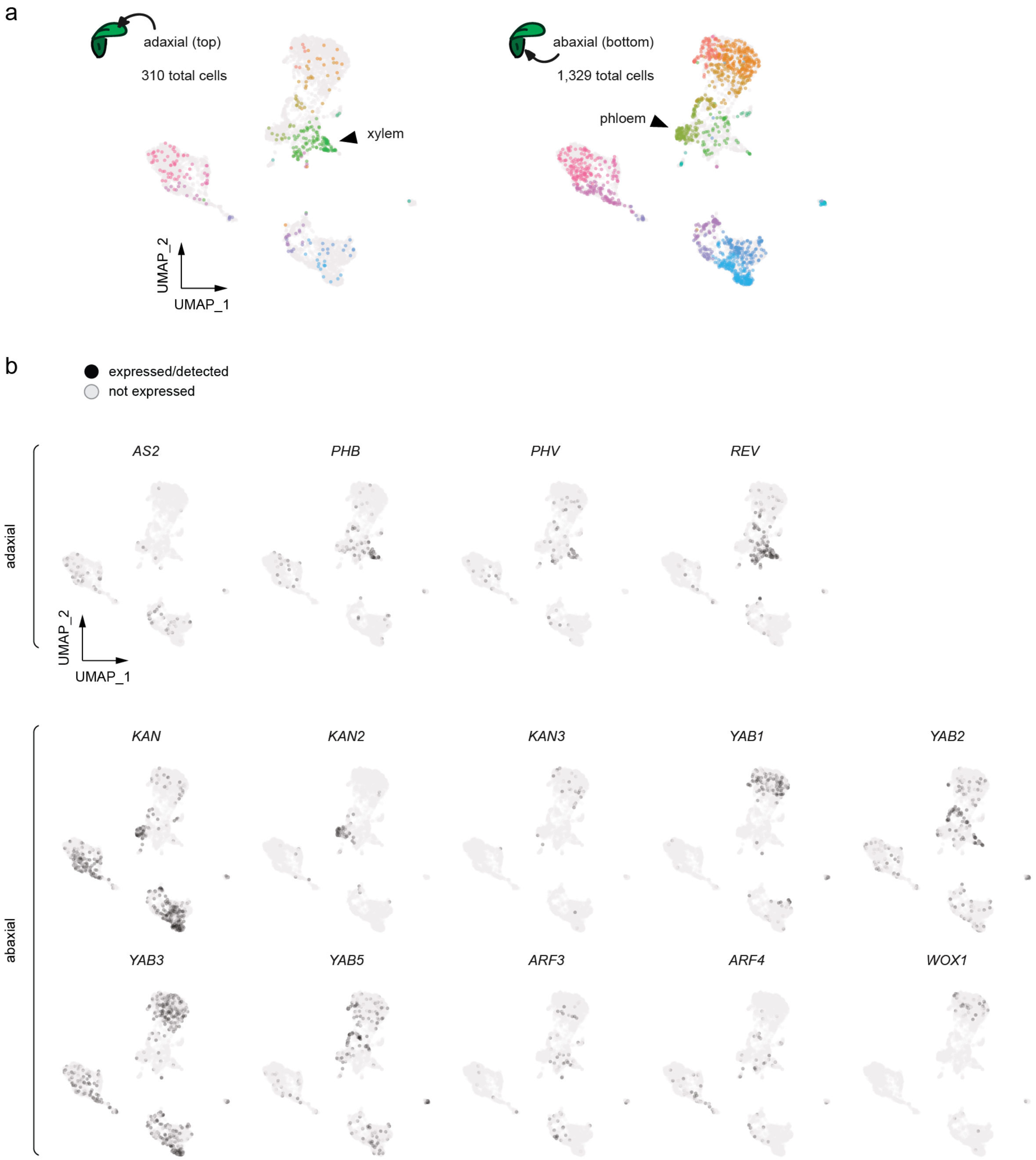


**Figure S4. Polarity signatures within the leaf atlas.** (a) UMAPs highlight cells (color corresponds to cluster resolution 1) with putative adaxial (top of leaf) versus abaxial (bottom) cell signatures. 310 and 1,329 total cells were scored as either adaxial or abaxial, respectively, based on whether they expressed at least one polarity marker (adaxial: *AS2*, *PHB*, *PHV*, *REV*; abaxial: *KAN*, *KAN2*, *KAN3*, *YAB1*, *YAB2*, *YAB3*, *YAB5*, *ARF3*, *ARF4*, *WOX1*). Abaxial markers *CRC*, *YAB4*, and *WOX3* were not expressed/detected. Not shown here, only 86 cells scored as 'ambiguous' since they co-expressed at least one adaxial and one abaxial marker. Note that among young guard cells (pink cluster) some fraction of cells still express polarity markers. (b) Expression profiles (black: expressed/detected, grey: not expressed/detected) of marker genes that define the leaf organ adaxial/abaxial polarity axis.

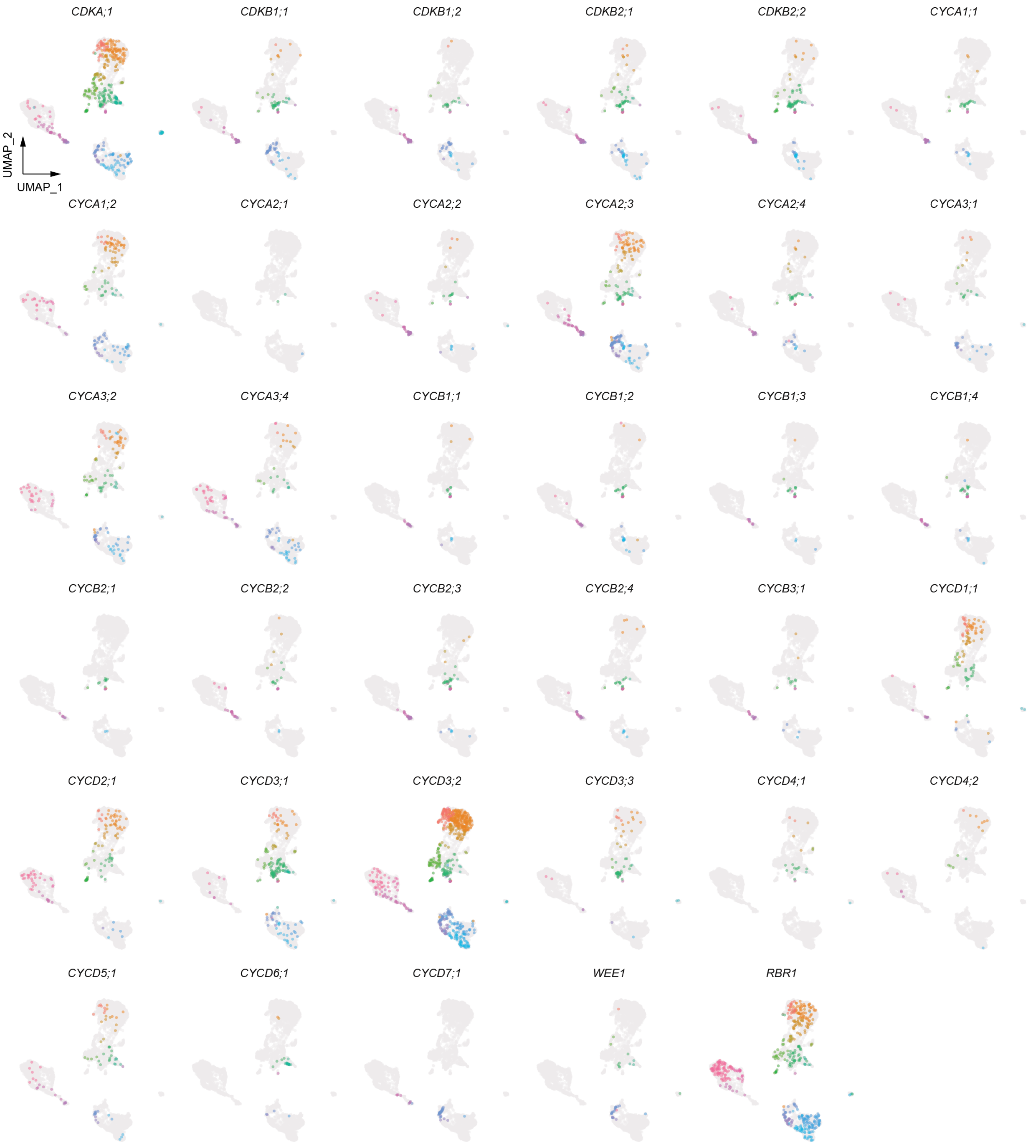


**Figure S5**. **Distinct and overlapping cell cycle signatures.** UMAPs of the developing leaf atlas highlight expression profiles (color corresponds to cluster resolution 1) of representative core cell cycle regulators. Shown here is a more complete set of regulators, which also includes those presented in Figure 3 for comparison.

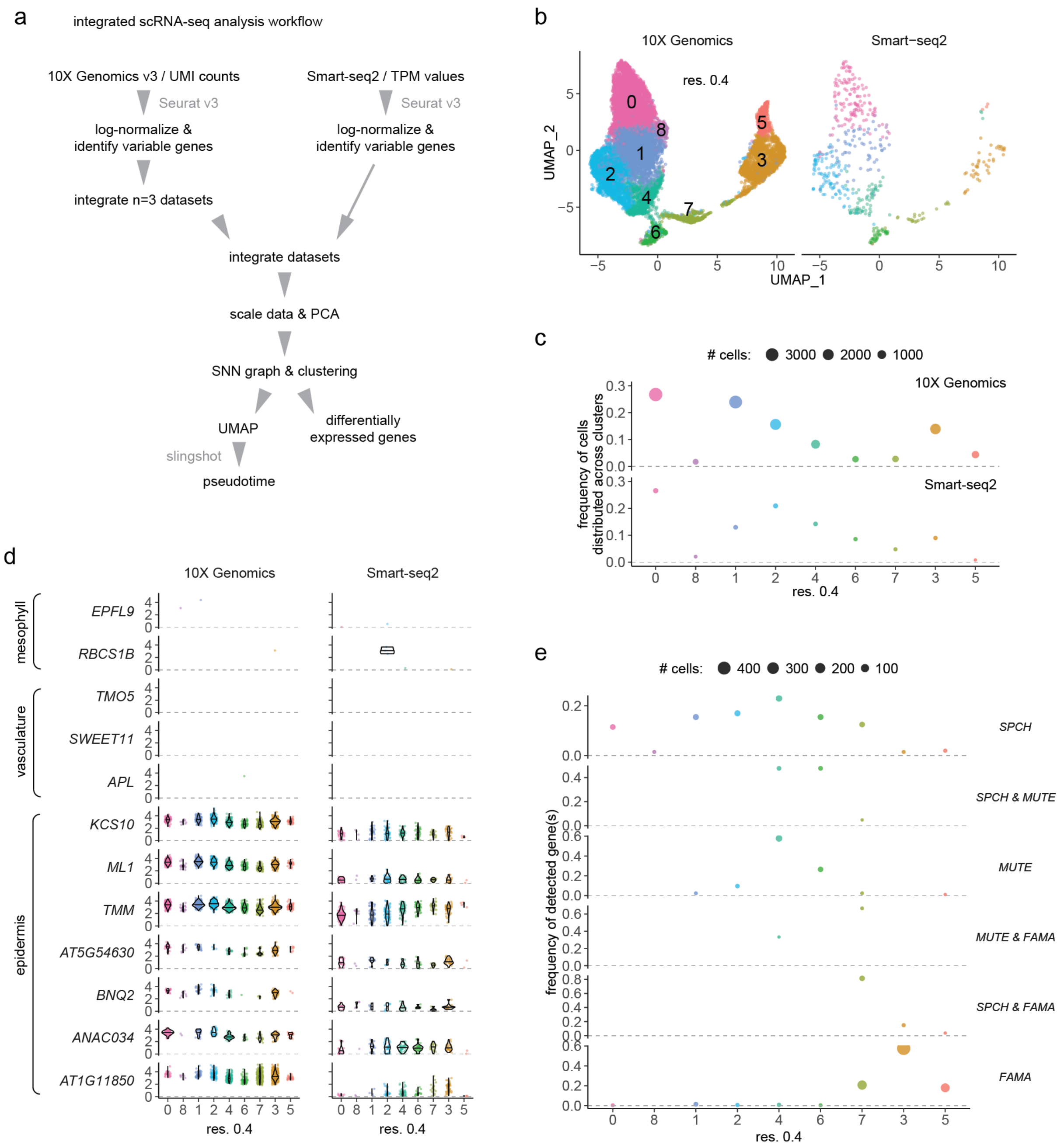


**Figure S6. Experimental workflow and dataset attributes for stomatal lineage profiles.** (a) Workflow used to integrate the stomatal lineage 10X Genomics and Smart-seq2 datasets from developing shoot tissue of 10 dpg *TMMp::TMM-YFP* seedlings. (b) Separate UMAPs of integrated datasets, along with (c) the frequency and number of cells from each dataset that contribute to a given cluster. (d) Violin plots show expression of given genes within datasets and clusters. Included are expression profiles of known regulators across epidermal, vasculature, and mesophyll leaf tissue. *AT5G54630*, *BNQ2*, *ANAC034*, and *AT1G11850* genes were found to be expressed in epidermal tissue in Figure 1. (e) Distribution of core transcriptional regulators (*SPCH*, *MUTE*, and *FAMA*) plotted as the frequency and number of cells that express a given gene combination within each cluster. Colors in (b-e) correspond to cluster resolution 0.4.

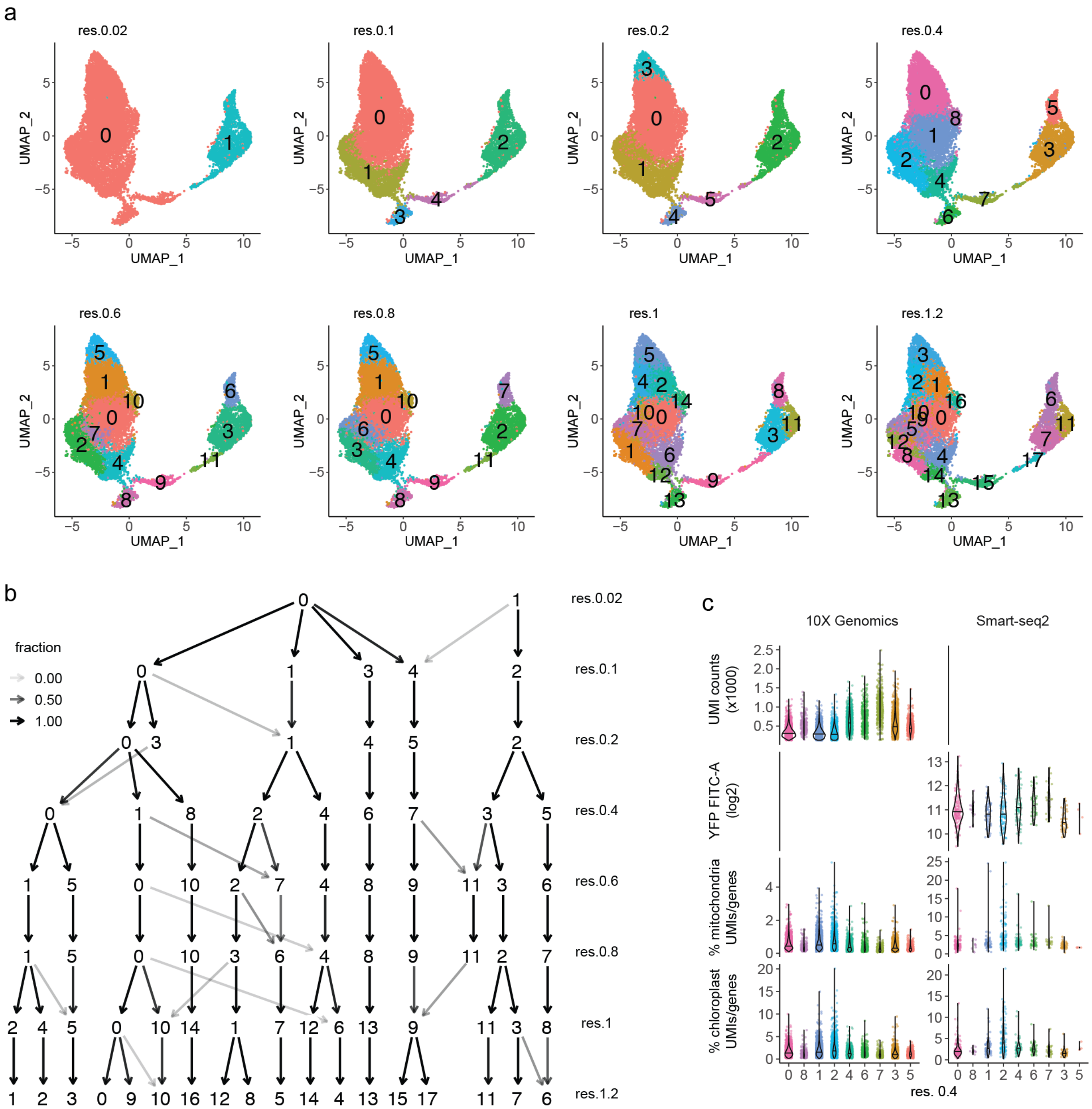


**Figure S7**. **Clustering analysis reveals similarities between clusters.** (a) Stomatal lineage UMAPs depict a range of clustering resolutions. (b) As described in Figure S2, clustering analysis with clustree shows how cells shift between clusters as the resolution is increased. (c) Additional quantification from respective datasets and clusters (color corresponds to cluster resolution 0.4).

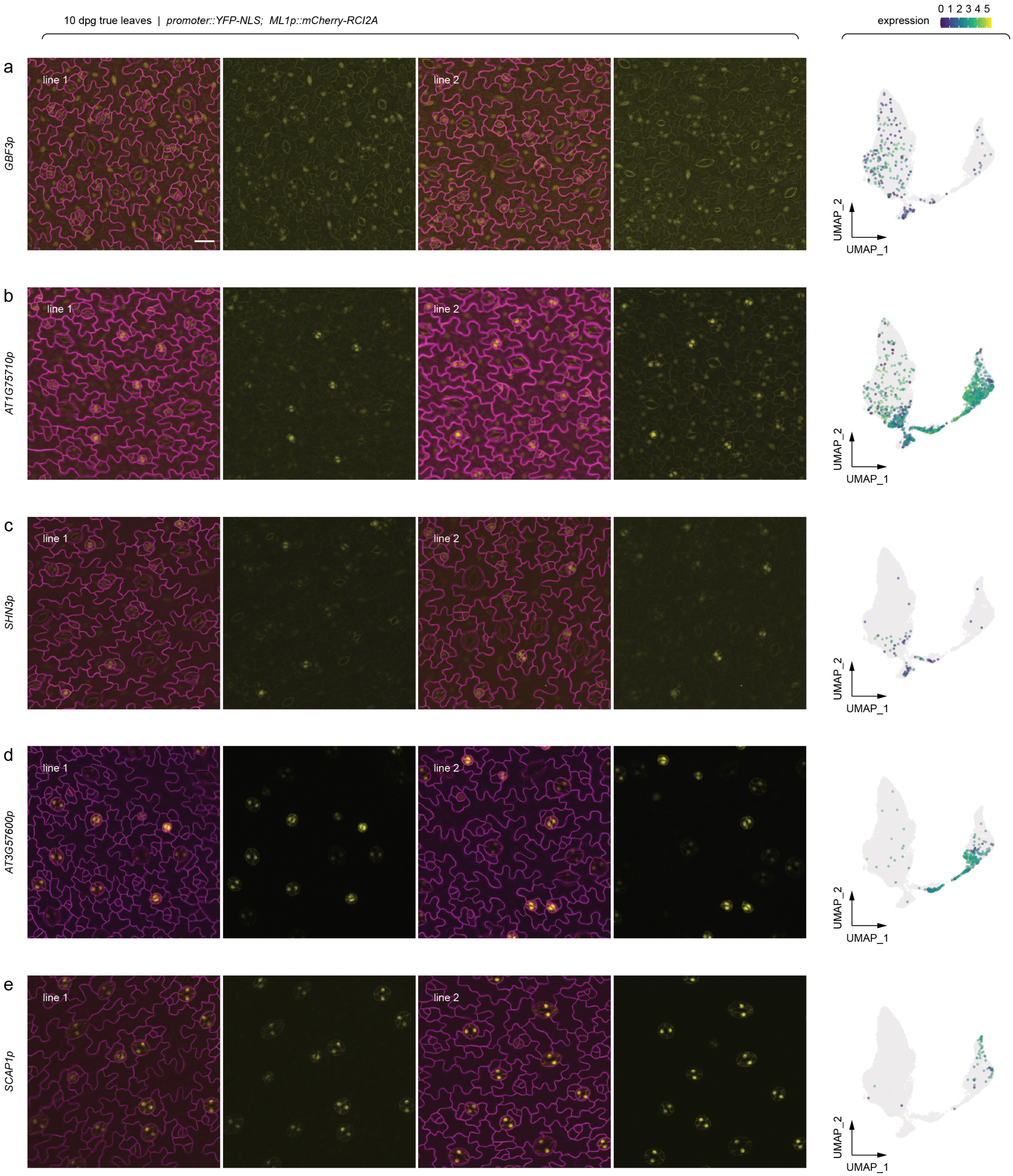


**Figure S8**. **Onset of predicted gene promoter activity and expression within the stomatal lineage.** As with Figure S3, shown are representative epidermal confocal images and corresponding scRNA-seq expression profiles. Representative confocal images of abaxial epidermis (first true leaves) are from 10 dpg seedlings of two independent reporter lines. Expression of *YFP-NLS* reporters were driven by respective promoters, and cell outlines were visualized by *ML1p::mCherry-RCI2A*. The same field with and without the *ML1p::mCherry-RCI2A* channel is presented for each reporter. Predicted scRNA-seq expression profiles are depicted in the corresponding UMAP plots on the far right. All images were taken at the same magnification (scale bar: 20 μm).

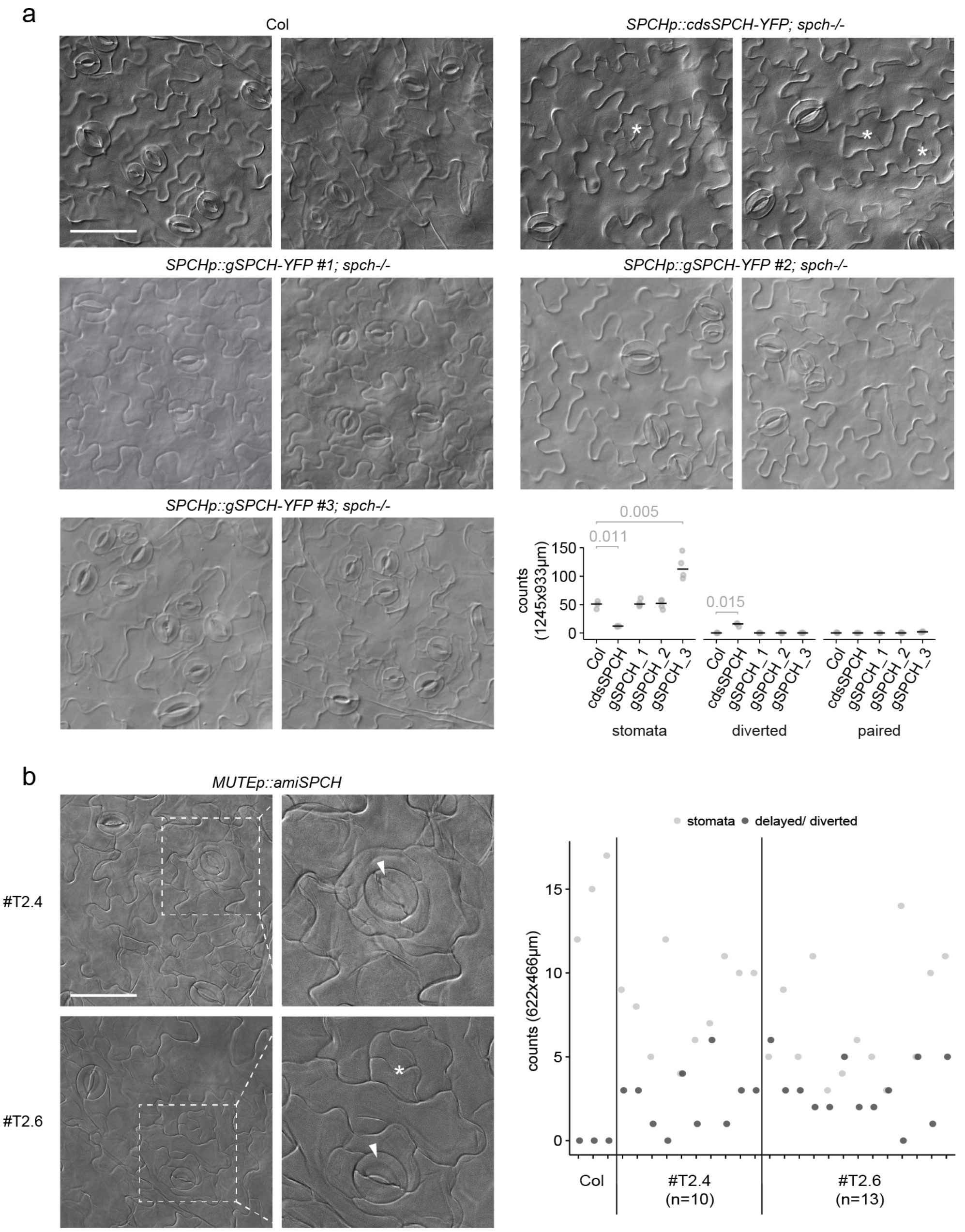


**Figure S9**. **The stomatal lineage requires fine-tuned *SPCH* gene expression.** (a) Cotyledon images and respective quantification from Col wildtype, *SPCHp::cdsSPCH-YFP spch-/-*, and *SPCHp::gSPCH-YFP spch-/-* . P-values are from the two-sample t-test to assess stomatal counts, whereas the one-sample t-test was used to assess diverted events. (b) Cotyledon images and respective quantification from two independent lines with the *MUTEp::amiSPCH* transgene. Representative DIC images were taken at 15 dpg, and the quantification includes stomata and diverted/delayed counts from individual images/seedlings. Plots include grey points that each represent counts from an individual seedling (at least n=3 seedlings were measured per genotype). All images were taken at the same magnification (scale bars: 100 µm), except for insets demarked by white dashed boxes. The asterisk indicates the border of diverted cell identities, and the arrowheads point to stomata at the center of putative diverted amplifying divisions.

### SUPPLEMENTARY TABLES

**Table S1.** Dataset attributes of scRNA-seq approaches; library preparation approaches, tissues and reporters sampled (from 10 dpg seedlings), total number of cells and mean values of reads that passed mapping preprocessing, and postprocessing median values of UMIs and genes.

**Table S2.** Primers used in the study to create reporter transgenes and lines expressing *amiRNAs*.

**Table S3.** Genes/AGI codes included in the study. TAIR (AGI, Arabidopsis Genome Initiative code), SYMBOL (gene symbol), and GENENAME annotations are from org.At.tair.db (Carlson, n.d.). Interpro_description annotations are from biomaRt (biomart = "plants_mart", host = "plants.ensembl.org", dataset = "athaliana_eg_gene") (Durinck et al., 2009, 2005).

**Table S4.** Cell cycle genes used for scoring and regression in Seurat v3 (Butler et al., 2018), with respective annotations as described in Table S3. These were identified as core regulators and enriched in previous time-course experiments (Menges et al., 2005, 2003; Vandepoele et al., 2002).

**Table S5.** Differentially expressed genes (a,e) (along with the top 50 per cluster, b,f) and transcription factors (c,g), as well as enriched GO (biological processes) (d,h), within the *ML1p::YFP-RCI2A* 10X Genomics leaf atlas. Included are the top enrichments the coarse resolution 0.1 (res. 0.1, a-d) and refined resolution 1 (res. 1, e-h) (Figure 1, Figure S2). Differentially expressed gene analysis; p_val: unadjusted p-value from Wilcoxon Rank Sum test, p_val_adj: adjusted p-value based on Bonferroni correction, avg_logFC: X-fold difference/change (log-scale) of the average expression between the two groups, FC: X-fold difference/change, pct.1: percentage of cells that express a gene within the given associated cluster (first group), pct.2: percentage of cells that express a gene across all the other clusters (second group), tf_family: transcription factor family. Enriched GO analysis; GO_ID: gene ontology ID, GO_description: gene ontology description, gene_ratio: ratio of genes detected within a given GO ID gene set, pvalue: p-value from hypergeometric test, p.adjust: adjusted p-value based on Bonferroni correction, qvalue: FDR adjusted p-value.

**Table S6.** Differentially expressed genes (a) and transcription factors (b) between adaxial (top) and abaxial (bottom) signatures (Figure S4) within the two main epidermal clusters of the *ML1p::YFP-RCI2A* 10X Genomics leaf atlas. 129 and 704 total cells were scored as either adaxial or abaxial, respectively, based on whether they expressed at least one polarity marker (adaxial: *AS2*, *PHB*, *PHV*, *REV*; abaxial: *KAN*, *KAN2*, *KAN3*, *YAB1*, *YAB2*, *YAB3*, *YAB5*, *ARF3*, *ARF4*, *WOX1*).

**Table S7.** Differentially expressed genes (a) (along with the top 50 per cluster, b) and transcription factors (c), as well as enriched GO (d), within *TMMp::TMM-YFP* integrated 10X Genomics and Smart-seq2 stomatal lineage cell states. Enrichments correspond to clusters from resolution 0.4 (Figure 4, Figure S7). Included are putative gene targets of SPCH, with annotated SPCH ChIP-seq binding events (Lau et al., 2014) and RNA-seq fold-changes detected in reprogrammed $FAMA^{LGK}$ cells (Lee et al., 2019), which reveals genes that correlate with induction of SPCH.
